# Sublethal and transgenerational effects of lambda-cyhalothrin and abamectin on the development and reproduction of *Cydia pomonella*

**DOI:** 10.1101/2022.07.05.498911

**Authors:** Di Ju, Yu-Xi Liu, Xue Liu, Youssef Dewer, David Mota-Sanchez, Xue-Qing Yang

## Abstract

The codling moth *Cydia pomonella* (Lepidoptera: Tortricidae) is a major invasive pest of pome fruits and walnuts worldwide. Lambda-cyhalothrin (LCT) and abamectin (AM) have been frequently used in *C. pomonella* control, but control of this pest is very difficult because shortly after hatching, larvae of this insect bore tunnels and hide inside host plant fruit. In this study, a simulated field spray bioassay method was developed against neonate larvae of *C. pomonella* and concentration-response bioassays were conducted to evaluate the susceptibility of the neonate larvae to LCT and AM. Exposure of neonate larvae to sublethal concentrations (LC_30_) of LCT or AM significantly reduced the survival rate of larvae (4th and 5th instars), lowered the mean weight of larvae and pupae, and decreased the daily maximal number of eggs laid and the total number of eggs laid (fecundity) per female. The sublethal effects, including reduced body mass, mean fecundity and net reproductive rate, extended mean generation time, and shortened oviposition period, were also found in transgenerational offspring. Furthermore, the transgenerational maternal effects were more obvious for AM than LCT, in comparison to the control. Additionally, the estimated population size was decreased by exposure to LC_30_ of LCT and AM, and the observed reduction of fecundity and population size within and across generations was likely the result of the downregulation of the reproduction-related vitellogenin gene (*CpVg*) after exposure to LC_30_ of LCT and AM. These results provide a better understanding of the overall effects of LCT and AM on *C. pomonella* and the transgenerational effects which should be taken into consideration when using insecticides in order to control *C. pomonella*.

## Introduction

Insecticide efficacy is often characterized by the mortality rate in pest populations within a given period. Nevertheless, apart from the strong and direct lethal effect induced by insecticides, sublethal effects likely occur following environmental degradation of insecticides or via heterogeneous spatial application on fields and orchards (Desneux et al., 2007; Ju et al., 2021). Besides mortality, the sublethal concentrations or doses of an insecticide can affect insect biology, physiology, behavior and demographic parameters, such as survival rate, developmental rate, longevity, fecundity, fertility, mating behavior, food searching and oviposition (Desneux et al., 2004; Dewer et al., 2016; Qu et al., 2017; Wang et al., 2018; Li et al., 2019; Khan et al., 2021., Mauduit et al., 2021; Wu et al., 2022). Moreover, stimulatory effects induced by low concentrations or doses of insecticides referred to as hermetic effects, may occur and have been observed in some species. For example, exposure of *Sitophilus zeamai*s to sublethal doses of pyrethroids resulted in a higher net reproductive rate (Guedes et al., 2010). Additionally, hormetic effects on the fitness traits of *Nilaparvata lugens* were observed, including the number of eggs laid per female and longevity of both susceptible and field-collected strains after exposure to their LC_20_ of nitenpyram for six generations (Gong et al., 2022). Therefore, the sublethal effects of insecticides may influence the population dynamics and outbreaks of pest species in generally unforeseen ways.

Additionally, effects induced by sublethal concentrations and doses of insecticides can be observed in transgenerational offspring, even when the offspring generation grows in favorable environments and without insecticides (Ayyanath et al., 2013; Baena-Díaz et al., 2018; Margus, et al., 2019; Gong et al., 2022). For example, exposure of parental generations of *Leptinotarsa decemlineata* to deltamethrin has negative influences on the offspring in terms of decreased body mass (Piiroinen et al., 2014). Previous research has suggested that insecticide exposure not only impairs the reproductive output of the exposed parental generation, but can also reduce reproduction in the offspring (Müller et al., 2017 & 2019). Another study investigated the fitness consequences of ivermectin residues in cow dung among adult *Sepsis punctum* and documented decreased offspring production when one or both of the parents were exposed to ivermectin, implying a reduction in fertility or fecundity (Conforti et al., 2018). Therefore, it is critical to understand the transgenerational effects of insecticides on pests in order to optimize the application of insecticides into integrated pest management (IPM) programs.

The codling moth *Cydia pomonella* (Lepidoptera: Tortricidae) is a major pest of pome fruits and walnuts worldwide (Yang & Zhang, 2015; Yang et al., 2016). The current estimated crop loss due to this pest is approximately US$10 million annually (Ju et al., 2021). Neonate larvae are the main targets of insecticides for controlling *C. pomonella* (Yang et al., 2016), but this pest is very difficult to control because shortly after neonate larvae bore tunnels and hide inside host plant fruit (Ju et al., 2022). Currently, *C. pomonella* control relies primarily on chemical insecticides such as lambda-cyhalothrin (LCT) and abamectin (AM), which are considered to be the most widely used insecticides (Reyes et al., 2007; Hu et al., 2020). In many countries including China, LCT and AM are the two most used insecticides for controlling *C. pomonella* (Wang et al., 2019; Hu et al., 2020; Ju et al., 2022). Although sublethal effects of LCT and AM on development and reproduction have been widely reported in pests of food crops and vegetables (Yin et al., 2008; Song & Zhang, 2013), their potential sublethal effects on fruit pests such as *C. pomonella* have not been previously documented.

This study developed a simulated field spray bioassay method against neonate larvae of this pest in order to determine whether transgenerational effects on the fecundity and fitness of *C. pomonella* were induced by exposures to sublethal concentrations of LCT and AM. We determined the toxicity of LCT and AM on neonate larvae of an insecticide susceptible strain and investigated the effects of LC_30_ of insecticides on reproduction and fitness-related traits of *C. pomonella* and their transgenerational effects. Effects of sublethal concentration of LCT and AM on the expression levels of a reproduction-related vitellogenin receptor (*CpVg*) were also investigated. These results can help improve understanding of the sublethal effects of insecticides on insects in general, and facilitate developing of effective control strategies that take into account these sublethal effects in IPM programs targeting of *C. pomonella*.

## 2 Method and Materials

### 2.1. Insects and chemicals

A susceptible strain of was maintained under laboratory conditions of 26°C ± 1°C, 60% ± 5% relative humidity (RH), and a 16:8 h (L:D) photoperiod for more than 60 generations (F 60) without exposure to any insecticides. A detailed description of the feeding method has been previously described by Wang *et al*. (2019). The technical grade of LCT and AM (purity >98%) was purchased from Aladdin Reagent (Shanghai, China).

### 2.2. Insecticide bioassays

To determine the log concentration-mortality response of neonate larvae to LCT, a stock solution (technical grade with purity >98% was purchased from Aladdin Reagent, Shanghai, China) was prepared in acetone and then diluted to six concentrations (2, 1, 0.5, 0.25, 0,125, 0.0625, 0 mg/L). Twenty newly hatched larvae (less than 24-h-old) were gently placed on a filter paper (Whatman No. 1) in a petri dish with a diameter of 9 cm, and each concentration was sprayed directly on the larvae (2-3 times) using an atomizer (fine drops of spray) at 10-20 cm from the petri dish (Figure 1A). After the insecticide solution was dried, larvae were kept individualized in glass tubes (2 × 5 cm) and fed with an artificial diet according to Brinton et al. (1969). Larvae treated with acetone only were set as control. Three replicates were carried out for each concentration treatment of each insecticide. The mortality was recorded at 24, 48, and 72 hours post exposure (hpe) to the treatments. Larvae were considered dead when no physical response was detected using an ink brush tool.

**Figure 1.**
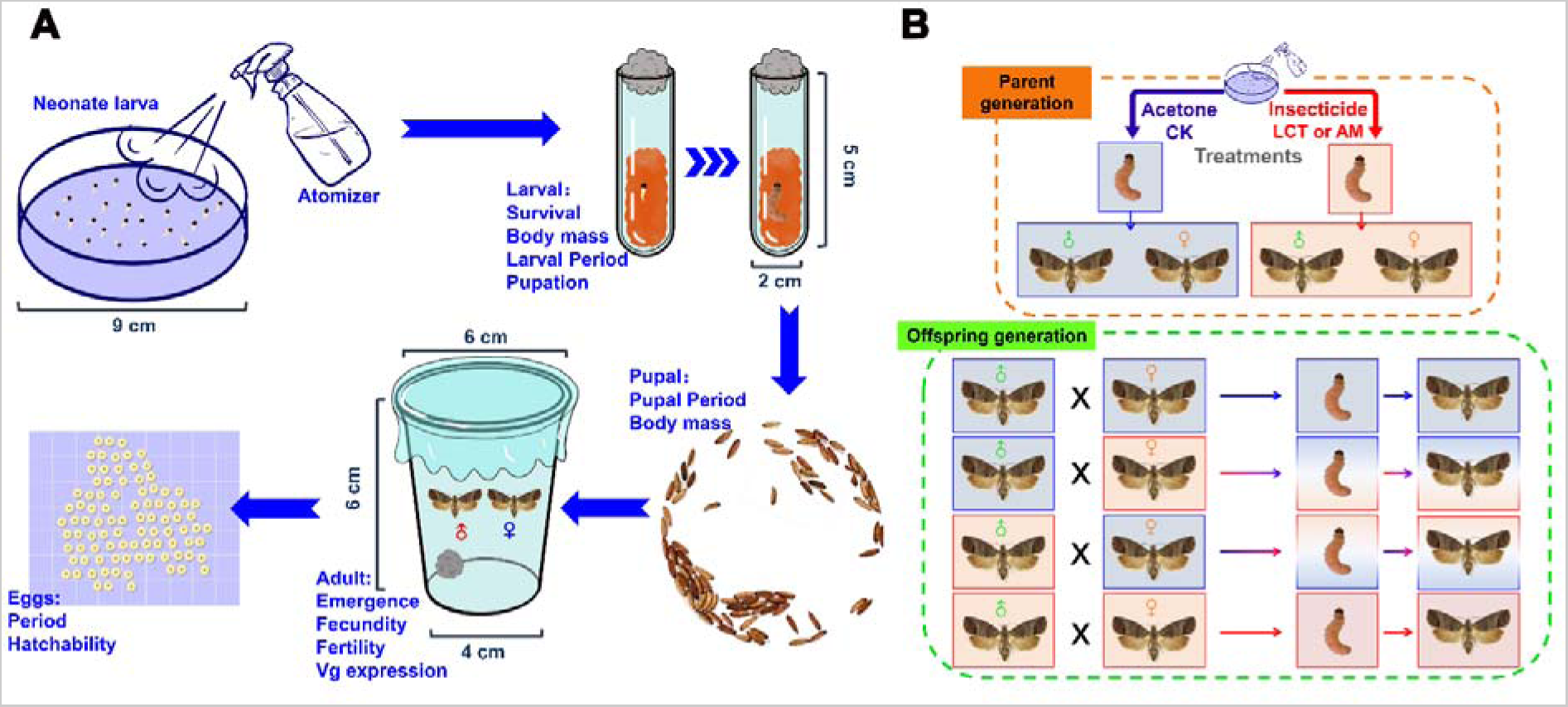
Schematic diagram of experimental design. (A) Schematic diagram of a simulated field spray bioassay method against neonate larvae, and rearing of *C. pomonella*. Twenty newly hatched larvae were gently placed on a filter paper (Whatman No. 1) in a petri dish with a diameter of 9 cm, and each concentration was sprayed directly on the larvae (2-3 times) using an automizer (fine drops of spray) at a distance of 10-20 cm from the petri dish. Larvae were kept individualized in glass tubes (2 × 5 cm) and fed with an artificial diet until pupation. Five pairs of adult moths were placed in a plastic cup container (6 × 4 × 6 cm), a piece of absorbent cotton soaked with a 10% honey solution was provided for adult feeding. (B) Schematic diagram of experimental design for sublethal, including two successive generations.The LC30 value of LCT and AM at 24 and 36 hours post exposure (hpe) were used as the exposure concentrations for sublethal effect experiments on *C. pomonella*.

The same method was used to determine the concentration-mortality of AM to neonate larvae at 36, 48, 72, and 96 hours due to slow kill activity of AM

### 2.3 Determination of lethal and sublethal concentrations

Mortality data were subjected to Probit analysis after correction of mortality (Abbott 1925). If the control mortality was higher than 5%, the mortality in the treatment group was corrected using Abbott’s formula. The concentrations of LCT and AM that would kill 10% (LC_10_), 30% (LC_30_), 50% (LC_50_), and 90% (LC_90_) of the individuals at 24 and 36 hpe were determined. To evaluate the sublethal effect of LCT and AM on *C. pomonella*, the LC_30_ concentration values of the two insecticides which were estimated through the concentration-mortality response mentioned was selected for the subsequent experiment.

### 2.4 Insecticide exposure and sublethal effects on the parent generation

Using the simulated field spray bioassay method as described above, neonate larvae were separately exposed to LC_30_ of LCT, AM and acetone. Three replicates, each with 150 larvae, were conducted for each concentration treatment level of each insecticide. The life stage and mortality of each instar were recorded from the third instar until all larvae either died or pupated. Body mass was measured at the third instar. In order to calculate pupation rates, larvae were placed into a plastic case (17 cm long, 8.5 cm wide, and 6.3 cm high) with soft paper on the bottom, and pupation was recorded daily. Pupae were sexed and weighed on the first day of pupation. The time and rate of adult eclosion were also recorded.

Female and male moths that emerged on the same day were randomly placed in a plastic cup container (6 cm top diameter, 4 cm bottom diameter, and 6 cm depth) with tiny air holes, and sealed with polyethylene plastic wrap. A piece of absorbent cotton soaked with a 10% honey solution was provided for adult feeding. Four groups were created including 1) control males paired with control females (CK♂ × CK♀); 2) insecticide treatment males paired with insecticide treatment females (LCT♂ × LCT♀ or AM♂ × AM♀); 3) control males paired with insecticide treatment females (CK♂ × LCT♀ or CK♂ × AM♀); 4) insecticide treatment males paired with control females (CK♀ × LCT♂ or CK♀ × AM♂). Five pairs were randomly introduced into an plastic container and were considered as a single replicate, and each treatment combination contained 15 replicates. The survival, number of eggs laid (fecundity) and hatched (fertility) of each adult were recorded daily until all the adults died. A schematic diagram of the experimental design is shown in Figure 1B.

On the 3rd, 4th, and 5th day following adult emergence (which represents the oviposition period from initial to peak), ten females from each repeating group were sampled and flash-frozen in liquid nitrogen and stored at −80 °C for later RNA extraction.

### 2.5 Sublethal effects on the traits of offspring

To evaluate the possible carryover effects of sublethal concentrations of LCT and AM on offspring, all eggs laid by adults from each treatment previously described were collected and the hatching rate for each group was recorded. Three replicates, each with 150 larvae hatched from each treatment, were supplied with a fresh untreated artificial diet until pupation. Larval and pupal developmental periods, body mass, mortality, pupation rate, pupal weight, adult eclosion rates, eclosion time, and fecundity were calculated using the same as method used in the parent generation.

### 2.6 Life table data analysis

The survival rate (*s_xj_*) (*x* = age, *j* = stage), which is the probability that a newly laid egg will survive to age *x* and stage *j*; and fecundity (*f_xj_*), which is the number of hatched eggs produced by a female at age *x* were calculated. The age-specific survival rate (*l_x_*) was then calculated as:

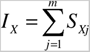

The net reproductive rate (*R_0_*) was calculated as:

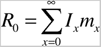

The intrinsic rate of increase (*r*) was calculated as:

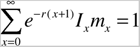

The mean generation time (*T*) in days was calculated as:

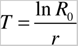

The raw life history data of individual *C. pomonella* were analyzed based on the age-stage, two-sex life table theory, and utilized the method described by Chi (Chi et al., 1988).

### 2.7 Population projection

The life table data for two generations following different treatments were used to project the population using the free TIMING-MSChart software program (Chi 2020), and this software program is available for download from http://140.120.197.173/Ecology/prod02.htm. The population size was projected over 120 days from an initial population.

### 2.8 Real-time quantitative PCR (RT-qPCR)

Total RNA was extracted from the whole bodies of a pool of 15 AM-treated larvae using the TaKaRa MiniBEST Universal RNA Extraction Kit (TaKaRa, Dalian, China) following the manufacturer’s instructions. The concentration and integrity of the RNA samples were determined using a NanoDrop2000 spectrophotometer (Thermo Fisher Scientific, Waltham, MA, USA) and a Agilent 2100 bioanalyzer (Agilent Technologies, Santa Clara, CA, USA), respectively. The RNA samples that met the requirements of an OD_260/280_ value of ≥1.8 and an integrity value of ≥ 7.0 were used for additional analyses.

The cDNA was synthesized using 1 µg of RNA from each treatment using the PrimeScript™ RT reagent Kit with gDNA Eraser (TaKaRa Dalian, China). RT-qPCR was performed on a Bio-Rad CFX96 (Bio-Rad, Hercules, CA USA) in a 20 µL reaction consisting of 10 μL of TB Green Premix Ex Taq 2 (TliRNaseH Plus; TaKaRa, Dalian, China), 1 µL each of forward and reverse primer, 1 µL of cDNA template, and 7 µL ddH_2_O. The RT-qPCR cycling was performed under the following conditions: 95°C for 30 s, followed by 40 cycles of 95°C for 5 s and 60°C for 30 s. A melting curve was conducted with a temperature increase from 63°C to 95°C with a ramp rate of 0.1°C/s to determine the specificity of the PCR products.

Using the vitellogenin mRNA sequence from *Spodoptera exigua* (GenBank: KT599434.1) as a query sequence, the *C. pomonella* vitellogenin gene (*CpVg*) was obtained (Figure S1) using the BLAST database tool (http://blast.ncbi.nlm.nih.gov/Blast.cgi) against the transcriptome SRR14923516 (Bioproject PRJNA741592) (InsectBase: CPOM075400). Molecular weight (Mw) and theoretical isoelectric points (pI) of *C. pomonella* vitellogenin protein were calculated using ExPASy (http://web.expasy.org/compute_pi/). The protein domain of CpVg was predicted using the NCBI web tool (https://www.ncbi.nlm.nih.gov/Structure/cdd/wrpsb.cgi). A phylogenetic tree was constructed by the neighbor-joining method using MEGA 5.0 software (Molecular Evolutionary Genetics Analysis using Maximum Likelihood, Evolutionary Distance, and Maximum Parsimony Methods) with 1000 bootstrap replicates.

Gene-specific primers (Table S1) for *CpVg* were designed with the Primer Premier 6 software (PREMIER Biosoft International, San Francisco, CA, USA). The β*-Actin* (KC832921) and *EF-1*α (MN037793) of *C. pomonella* were used as reference genes (Wei et al., 2020). Negative controls with RNase-free water instead of cDNA templates were conducted to exclude any potential genomic DNA contamination. Each reaction was carried out with three technical replicates for each RNA sample. The amplification curves, melting peaks, and melting curves are shown in Figure S2. The relative expression level of *CpVg* was calculated using the 2^−△△CT^ method (Livak & Schmittgen, 2001).

### 2.9 Data Analysis

The life history traits including the survival rate, body mass, development duration, emergence rate, number of eggs laid (fecundity), and eggs hatched (fertility) of *C. pomonella* were subjected to a one-way analysis of variance (ANOVA) with Duncan’s test using SPSS 20 software (IBM corporation, Armonk, NY, USA). A significance induction of *CpVg* gene expression by sublethal concentrations of LCT and AM was performed by an independent-samples T-test (\**P* < 0.05; \*\**P* < 0.01; \*\*\**P* < 0.001) using SPSS 20 software. All data are presented as the mean of triplicates ± standard deviation (SD).

## 3 Results

### 3.1 Determination of sublethal concentrations

The toxicity of LCT and AM to *C. pomonella* neonate larvaewere determined using a simulated field spray bioassay method. The linear regression of concentration-mortality relationship was fitted to actual data for the two insecticides tested. The LC_10_, LC_30_, LC_50_, and LC_90_ values was estimated as 0.001 mg/L (lower and upper limit: 0.000 and 0.007 mg/L respectively), 0.025 mg/L (lower and upper limit: 0.002 and 0.064 mg/L respectively), 0.218 mg/L (lower and upper limit: 0.099 and 0.376 mg/L respectively), and 43.297 mg/L (lower and upper limit: 8.706 and 3154.035 mg/L respectively) at 24 hpe for LCT (Table 1), and 0.007 mg/L (lower and upper limit: 0.003 and 0.012 mg/L respectively), 0.028 mg/L (lower and upper limit: 0.018 and 0.040 mg/L respectively), 0.075 mg/L (lower and upper limit: 0.054 and 0.099 mg/L respectively), and 0.798 mg/L (lower and upper limit: 0.545 and 1.328 mg/L respectively) at 36 hpe for AM (Table 2). The LC_30_ value of LCT and AM at 24 and 36 hpe were used as the exposure concentrations for sublethal effect experiments on *C. pomonella*.

**Table 1.**
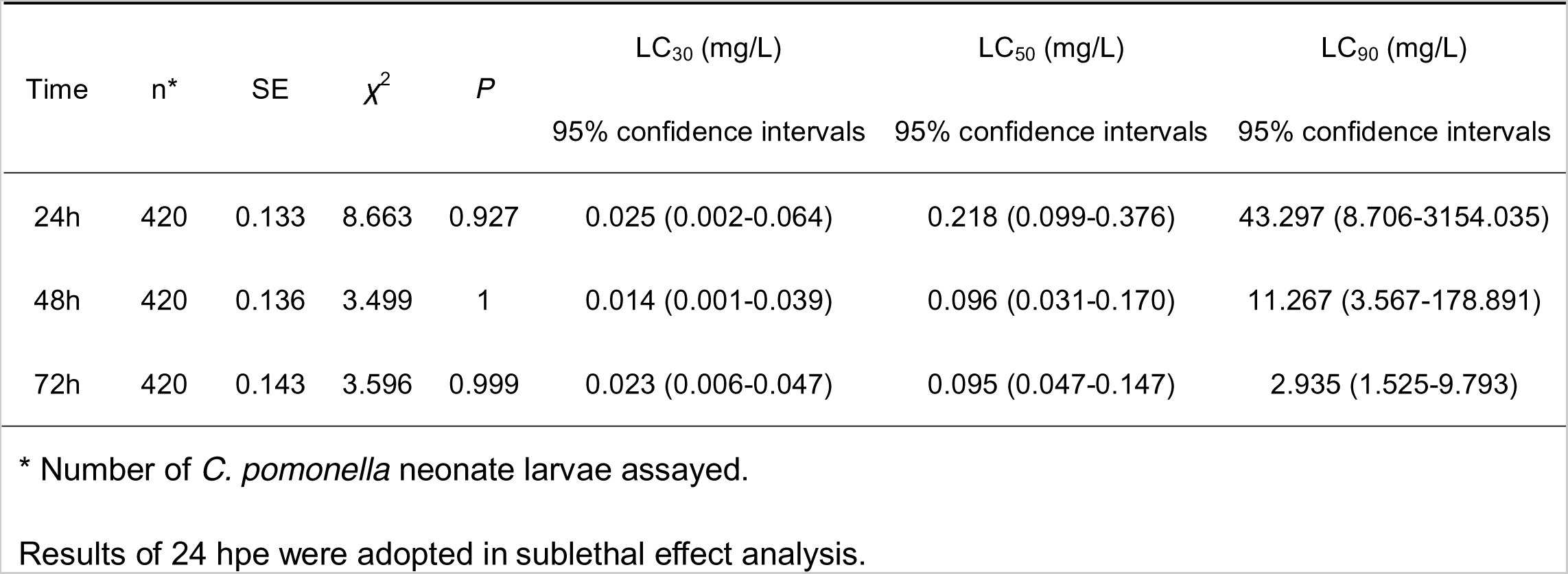
Relative toxicity of lambda-cyhalothrin (LCT) to the neonate larvae of Cydia pomonella.

| Time | n* | SE | $\chi^2$ | P | LC<sub>30</sub> (mg/L) | LC<sub>50</sub> (mg/L) | LC<sub>90</sub> (mg/L) |
| --- | --- | --- | --- | --- | --- | --- | --- |
|  |  |  |  |  | 95% confidence intervals | 95% confidence intervals | 95% confidence intervals |
| 24h | 420 | 0.133 | 8.663 | 0.927 | 0.025 (0.002-0.064) | 0.218 (0.099-0.376) | 43.297 (8.706-3154.035) |
| 48h | 420 | 0.136 | 3.499 | 1 | 0.014 (0.001-0.039) | 0.096 (0.031-0.170) | 11.267 (3.567-178.891) |
| 72h | 420 | 0.143 | 3.596 | 0.999 | 0.023 (0.006-0.047) | 0.095 (0.047-0.147) | 2.935 (1.525-9.793) |

\* Number of *C. pomonella* neonate larvae assayed.
Results of 24 hpe were adopted in sublethal effect analysis.

**Table 2.** Relative toxicity of abamectin (AM) to the neonate larvae of Cydia pomonella.

| Time | n* | SE | $\chi^2$ | P | LC<sub>10</sub> (mg/L) | LC<sub>30</sub> (mg/L) | LC<sub>50</sub> (mg/L) | LC<sub>90</sub> (mg/L) |
| --- | --- | --- | --- | --- | --- | --- | --- | --- |
|  |  |  |  |  | 95% confidence intervals | 95% confidence intervals | 95% confidence intervals | 95% confidence intervals |
| 36h | 450 | 0.118 | 12.091 | 0.986 | 0.007 (0.003-0.012) | 0.028 (0.018-0.040) | 0.075 (0.054-0.099) | 0.798 (0.545-1.328) |
| 48h | 450 | 0.120 | 8.143 | 0.999 | 0.004 (0.002-0.008) | 0.019 (0.011-0.029) | 0.053 (0.037-0.072) | 0.635 (0.431-1.073) |
| 72h | 450 | 0.132 | 10.61 | 0.995 | 0.005 (0.002-0.009) | 0.019 (0.011-0.028) | 0.048 (0.034-0.064) | 0.460 (0.321-0.746) |
| 96h | 450 | 0.135 | 10.274 | 0.996 | 0.004 (0.002-0.007) | 0.016 (0.009-0.023) | 0.040 (0.027-0.054) | 0.395 (0.276-0.642) |

\* Number of *Cydia pomonella* larvae assayed.
Results of 36 hpe were adopted in sublethal effect analysis.

### 3.2 Sublethal effects of LCT and AM on survival and development of two successive generations of *C. pomonella*

The survival rates of larvae, especially 4th (*l_4_*) and 5th (*l_5_*) instars were significantly decreased by treatment with LC_30_ of LCT and AM in the parental generation in comparison to the control. However, no significant difference in the survival rate of larvae was observed at offspring generation, compared with the control (Table 3).

**Table 3.**
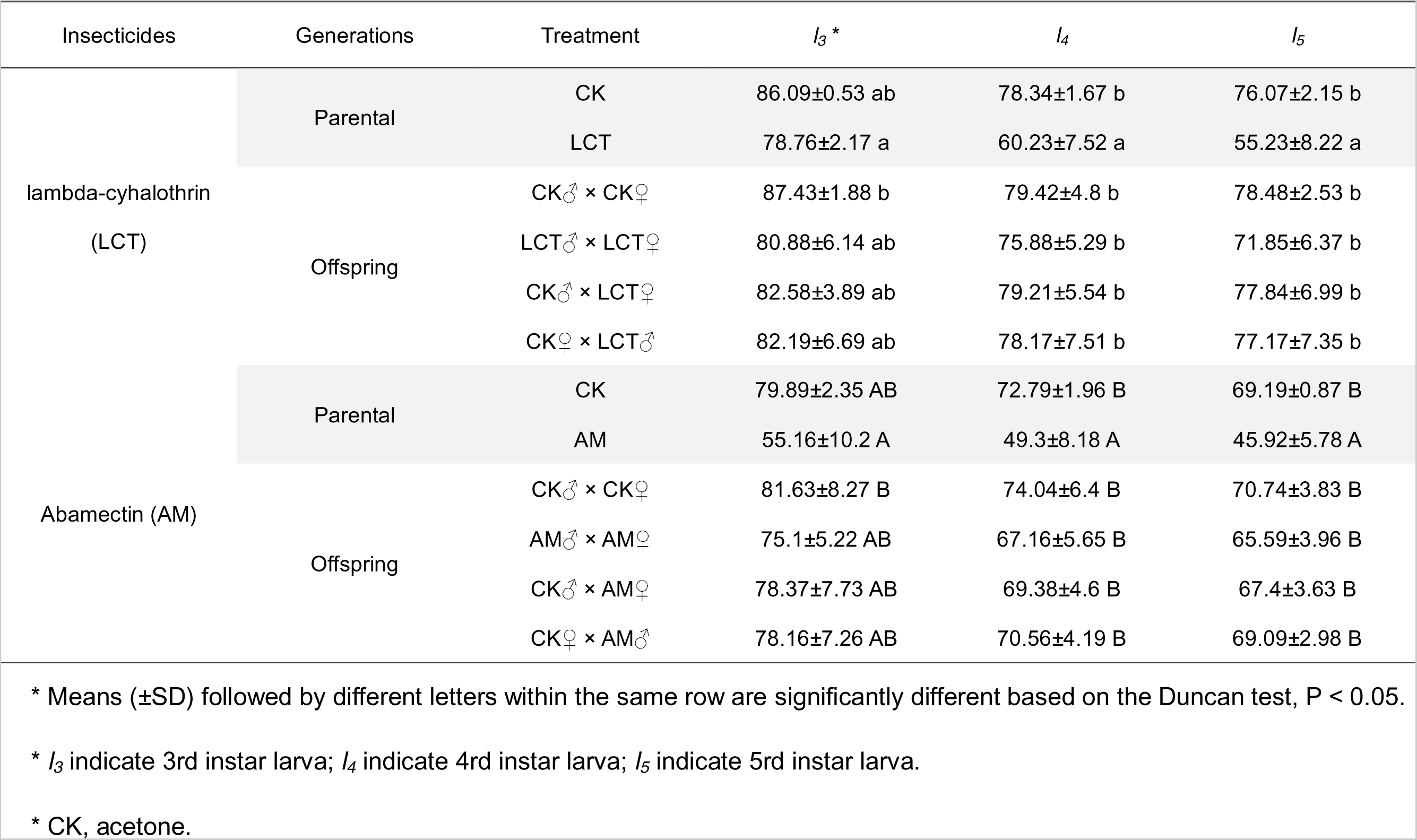
Sublethal effects of lambda-cyhalothrin (LCT) and abamectin (AM) on the survival of *Cydia pomonella* in two successive generations.

The body mass of 3^rd^ instar larvae (1.05 ± 0.11 mg) and female pupae (2.97 ± 0.08mg) in the parental generation was significantly reduced when neonate larvae were exposed to the LC_30_ concentration of LCT compared with the control. In the offspring generation, the body masses of larvae and pupae were still influenced by LCT treatment (Table 4). AM exposure resulted in lower larvae and male pupae weight in the parental generation, and the body mass of both larvae and pupae in the offspring generation in which females were treated, compared with the control (Table 4).

**Table 4.**
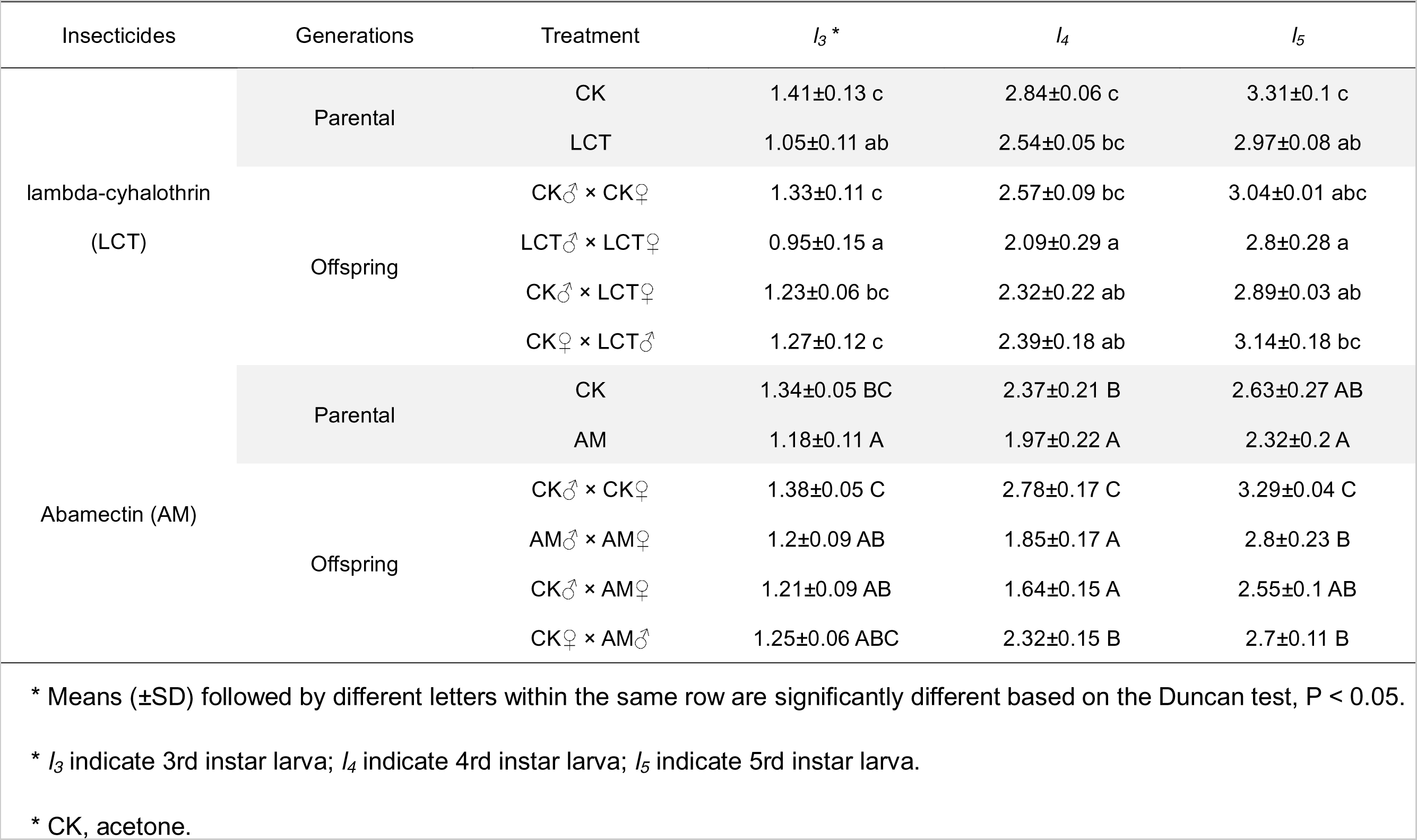
Sublethal Effects of lambda-cyhalothrin (LCT) and abamectin (AM) on larva and pupa weight of Cydia pomonella in two successive generations.

There was not statistically significant differences of both insecticides on the developmental period in the parental generation. However, when both parents were treated with LCT or AM, the offspring larval stage was prolonged (Table 5). It is noteworthy that the egg period of offspring produced by LCT treated females paired with non-treated males was prolonged (6.9 ± 0.66 days), compared with control (5.3 ± 0.82 days) (Table 5).

**Table 5.**
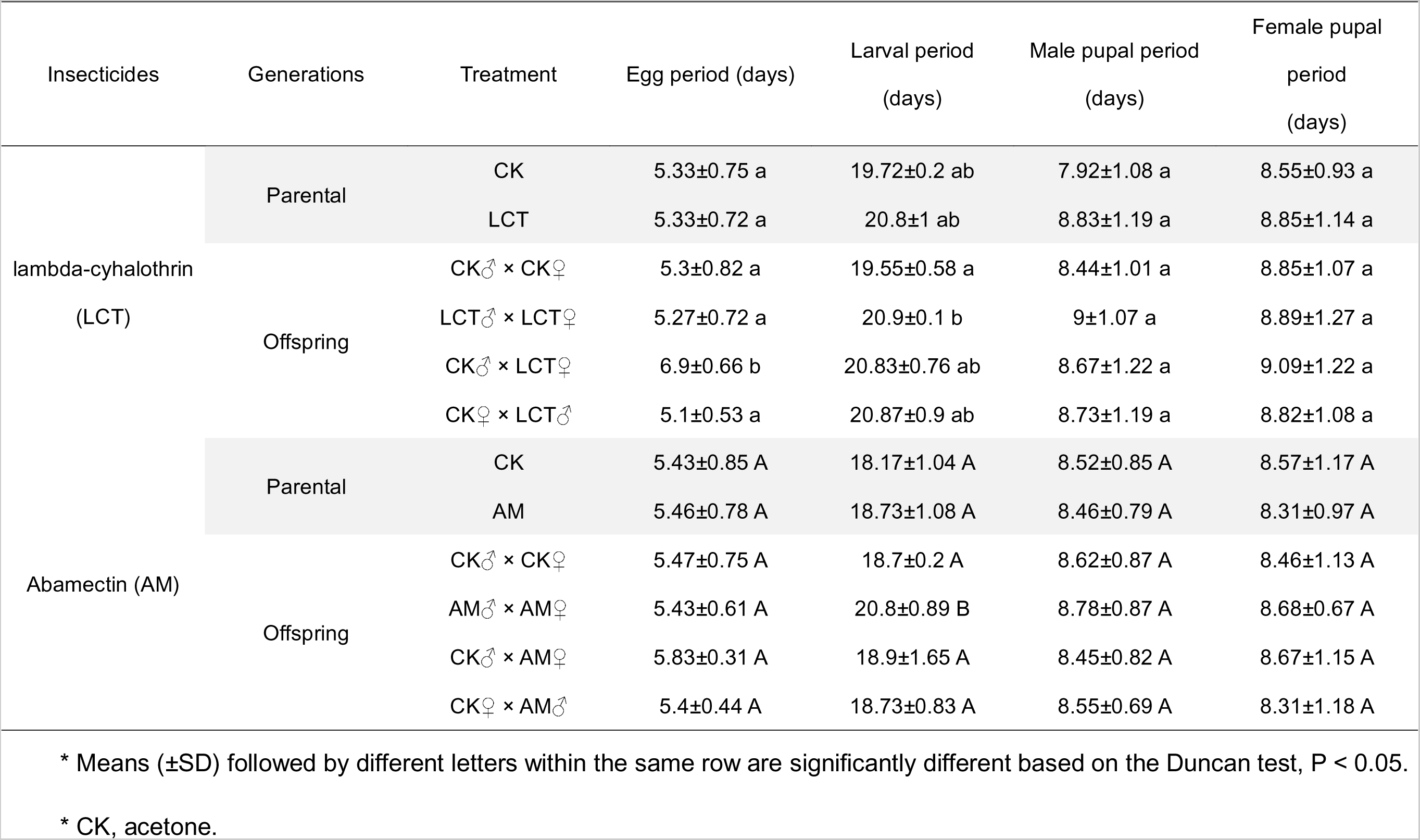
Sublethal effects of lambda-cyhalothrin (LCT) and abamectin (AM) on the development and reproduction of *Cydia pomonella* in two successive generations.

### 3.3 Sublethal effects of LCT and AM on pupation and adult emergence of two successive generations of *C. pomonella*

The pupation and emergence rates of two successive generations of *C. pomonella* were not affected by exposure to LC_30_ of both insecticides compared with the control (Figure S3).

### 3.4 Sublethal effects of LCT and AM on fecundity and fertility of two successive generations of *C. pomonella*

The total number of eggs laid (fecundity) and hatched (fertility) by each female were compared between the insecticide-treated groups and the control group (Figure 2 & Figure 3).

**Figure 2.**
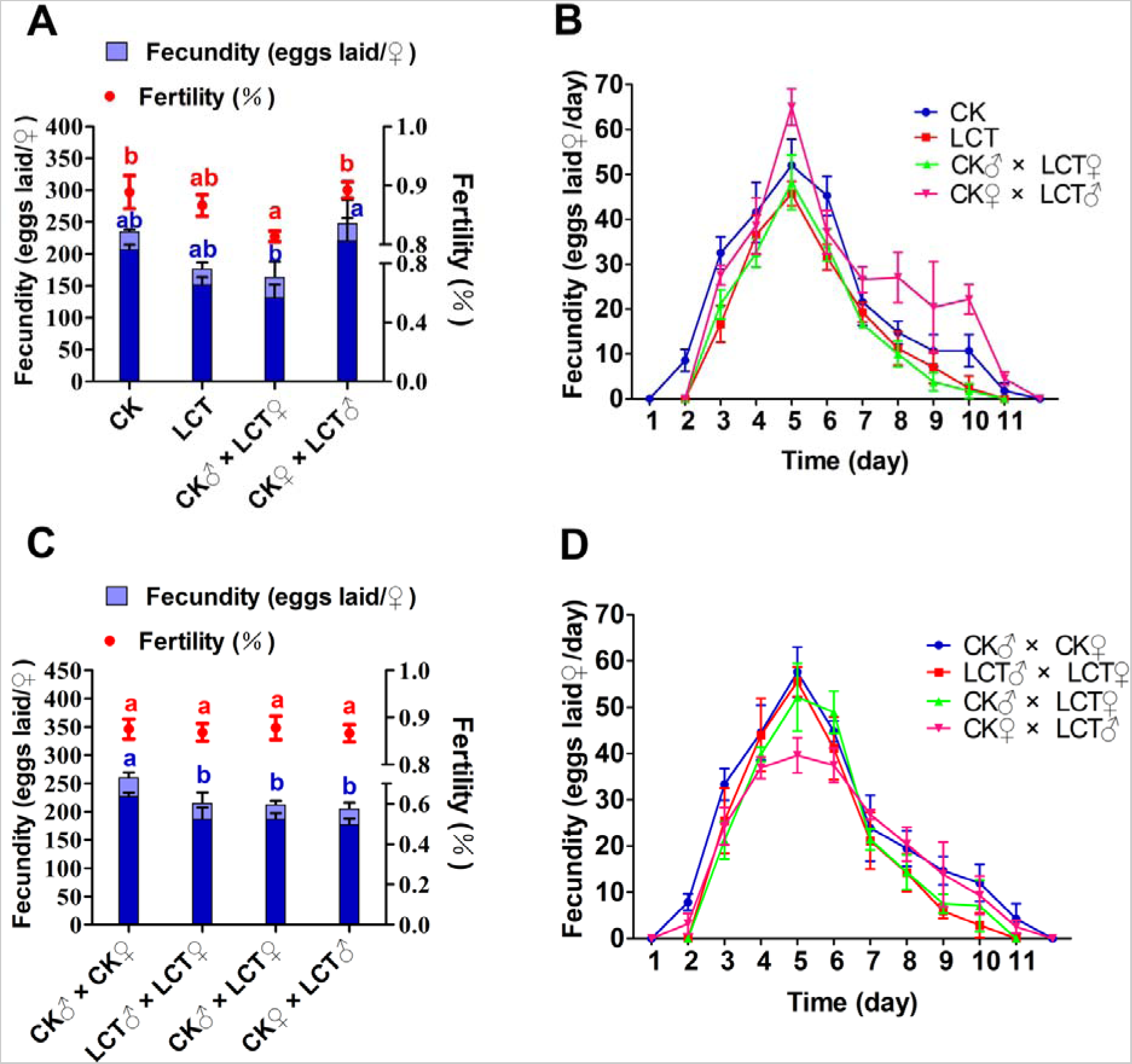
Sublethal effects of LCT on fecundity of *C. pomonella* adults. (A) The total number of eggs laid per female and hatchability of different treatments in the parental generation. (B) The daily number of eggs laid per female of different treatments in the parental generation. (C) The total number of eggs laid per female and hatchability of different treatments in the offspring generation. (D) The daily number of eggs laid per female of different treatments in the offspring generation. Light blue and dark blue columns represent the total number of eggs laid per female and the number of hatched larvae, respectively. Red dots represent the hatchability of eggs. CK, acetone; LCT, lambda-cyhalothrin. The results are shown as the mean ± SD. The error bars represent the standard errors calculated from three replicates. Letters on the error bars indicate significant differences analyzed by the one-way analysis of variance (ANOVA) with Duncan’s test (P < 0.05).

**Figure 3.**
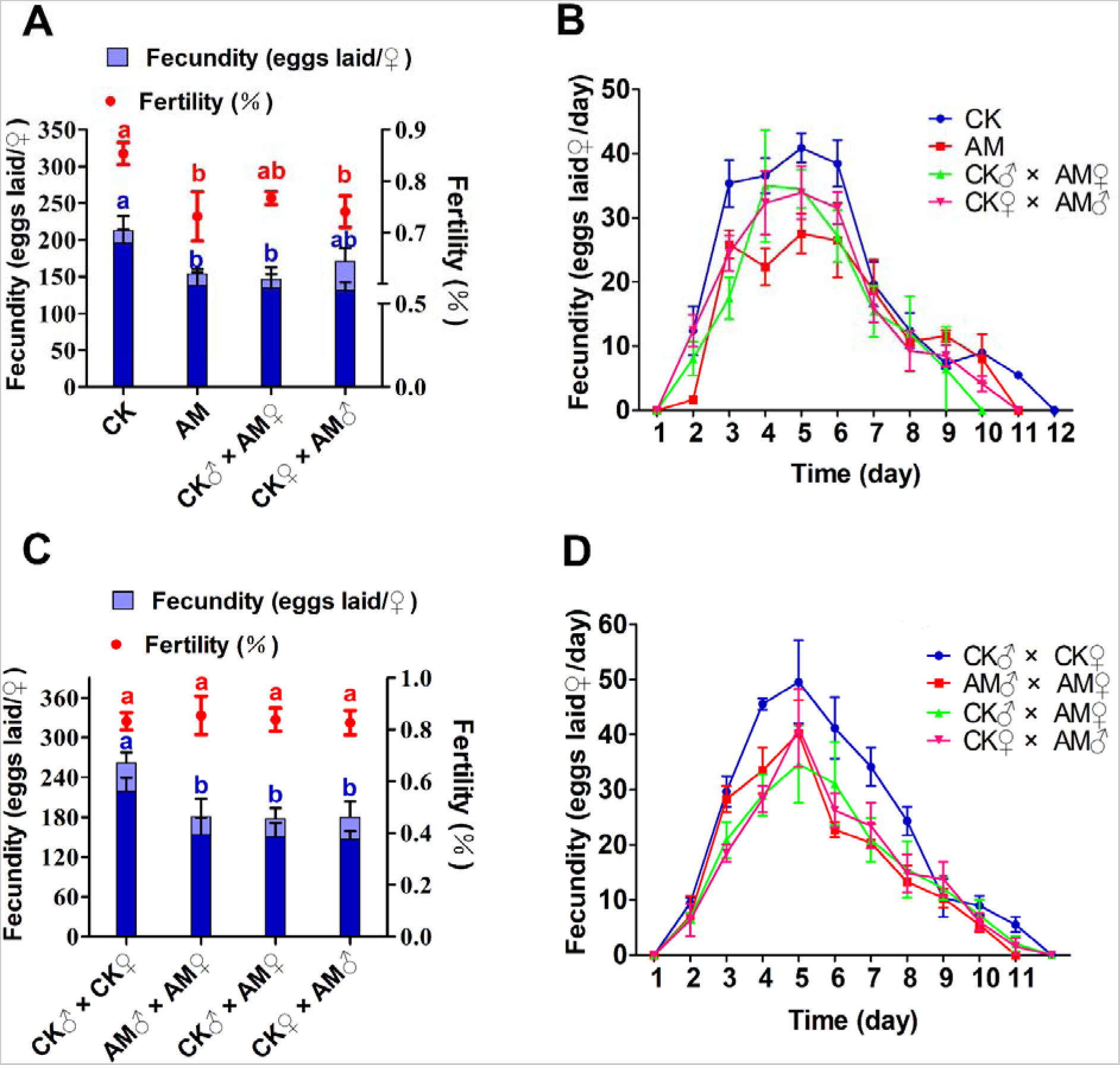
Sublethal effects of AM on fecundity of *C. pomonella* adults. (A) The total number of eggs laid (fecundity) and eggs hatched (fertility) exposed to LC_30_ of LCT and AM in the parental generation. (B) The daily number of eggs laid per female of different treatments in the parental generation. (C) The total number of eggs laid per female and hatchability of different treatments in the offspring generation. (D) The daily number of eggs laid per female of different treatments in the offspring generation. Light blue and dark blue columns represent the total number of eggs laid per female and the number of hatched larvae, respectively. Red dots represent the hatchability of eggs. CK, acetone; AM, abamectin. The results are shown as the mean ± SD. The error bars represent the standard errors calculated from three replicates. Letters on the error bars indicate significant differences analyzed by the one-way analysis of variance (ANOVA) with Duncan’s test (P < 0.05).

For the parental generation, after exposure of neonate larvae to LCT, the fecundity (Figure 2A), oviposition period, and the peak of eggs laid (Figure 2B) of the emerged females paired with non-treated males were decreased. Furthermore, the fertility of CK♂ paired with LCT♀ was significantly reduced compared with the control (Figure 2A). The fertility of LCT-treated groups was not suppressed in offspring (Figure 2C), but the fecundity and oviposition period were significantly decreased in the offspring when either of their parents were exposed to LCT, in comparison with the control (Figure 2C & 2D).

Exposure of neonate larvae to LC_30_ of AM was ween to significantly reduce the total number of eggs laid (Figure 3A), and the oviposition period and the peak of eggs laid (Figure 3B) when at the parental generation. The fecundity of females after copulation decreased when the mating partners (females or male, or both) were exposed to AM compared with the control (Figure 3C). No significant difference in the fertility between AM-treated groups and the control was found in the offspring (Figure 3C). However, a reduction in the oviposition period and the peak of eggs laid in AM-treated groups was also observed in offspring generation, compared with the control (Figure 3D).

### 3.4 Population parameters of two successive generations of *C. pomonella*

The population parameters in two successive generations of *C. pomonella* neonate larvae exposed to LC_30_ of LCT or AM were investigated. Results showed that neither the finite rate of increase (λ) nor the intrinsic rate (*r*) was significantly affected by the two insecticides (Table 6).

**Table 6.**
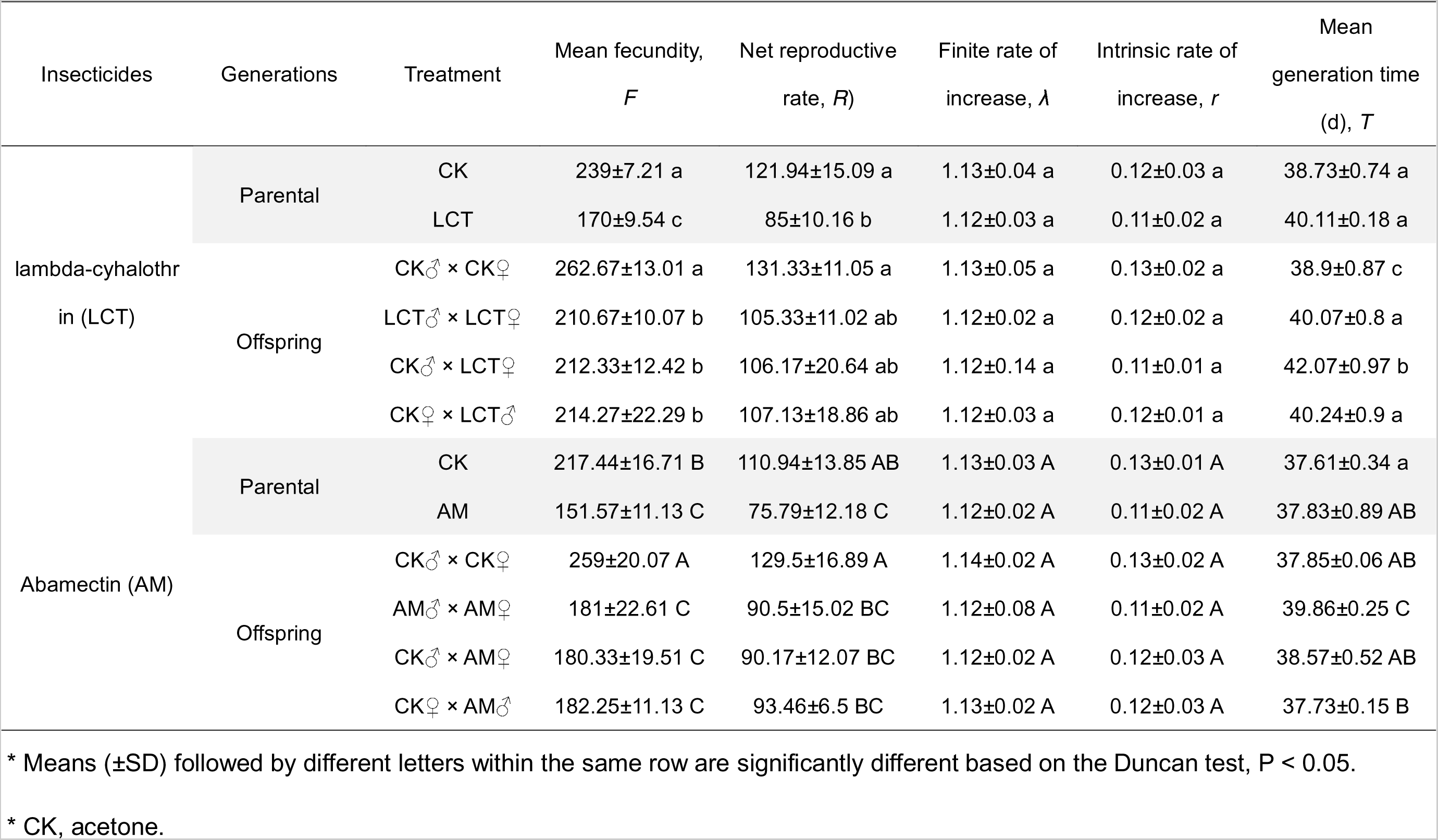
Sublethal effects of lambda-cyhalothrin (LCT) and abamectin (AM) on the population parameters of C. pomonella in two successive generations.

In *C. pomonella* neonate larvae exposed to LC_30_ of LCT, the *F* (mean fecundity), and *R_0_* (net reproductive rate) were both significantly decreased compared with the control in the parent generation, and the reduction in *F* persisted into the offspring. The *T* (mean generation time) of all pairs (LCT♂ × LCT♀, CK♂ × LCT♀, CK♀× LCT♂) in offspring (>40 days) was significantly increased when their parents were treated with LCT compared to the control (38.9 ± 0.87 days), although no significant prolongation of *T* was observed in the parent generation (Table 6).

Exposure of *C. pomonella* neonate larvae to LC_30_ of AM also reduced the *F* and *R_0_* in both parent and offspring compared to their corresponding controls. Interestingly, the *F* was reduced by 30% in both offspring and parents, in comparison to the controls (Table 6). This result indicated that *T* was significantly prolonged only in the offspring whose parents were both treated with AM during their neonate larva stage (Table 6).

Results of age-stage-specific survival analyses show that the *s_xj_* value for both male and female adults was not affected by LCT, whereas the value was lower in AM-treated populations compared with the control in the parental generation. The shortening of adult longevity was also observed in their descendants, especially in populations whose mothers were treated with AM at the neonate larvae stage (Figure 4).

**Figure 4.**
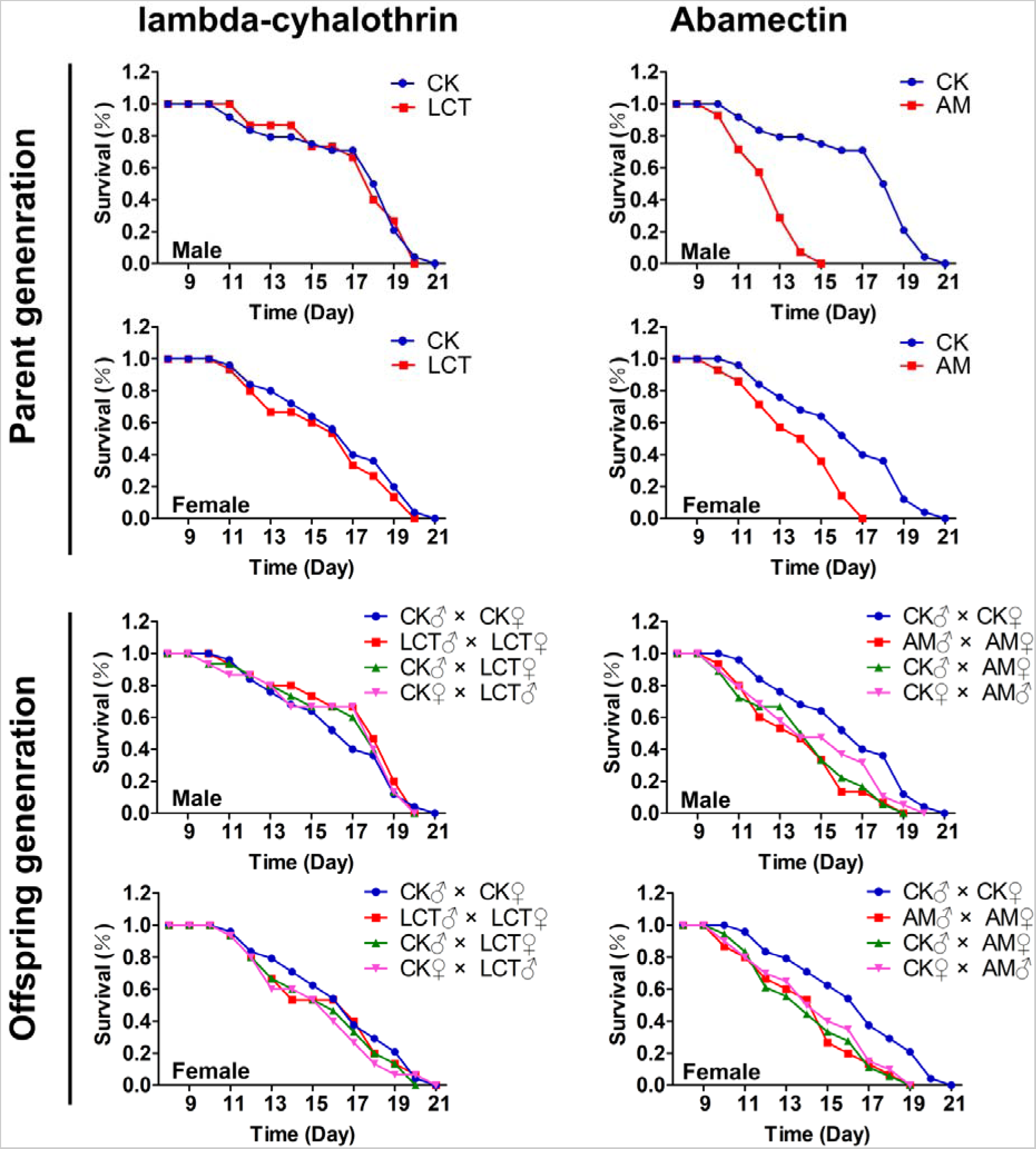
The survival rate of *C. pomonella* adults in different treatments. CK, acetone; LCT, lambda-cyhalothrin; AM, abamectin.

The age-stage-specific reproductive values (*v_xj_*) of *C. pomonella* represented the contribution of an individual at age *x* and stage *j* to the future population. The *v_xj_* value increased significantly when females began to produce viable eggs, and reached a peak of 160 in the control group, while in the LCT treatment group, the peak was less than 120 in the parental generation. For offspring, the *v_xj_* value increased at a delayed age (after the 30th day) and reached a peak value lower than that observed in the control. Similar to LCT observations, the highest *v_xj_*value seen in AM-treated groups was lower than that of the control in the parental generation. In the offspring generation, the increase in *v_xj_* value occurred earlier (at the age of 30 days) and remained at a high reproduction *v_xj_* value (>140) for a few days, however, a lower *v_xj_* value (<120) was observed in AM-treated groups (Figure 5).

**Figure 5.**
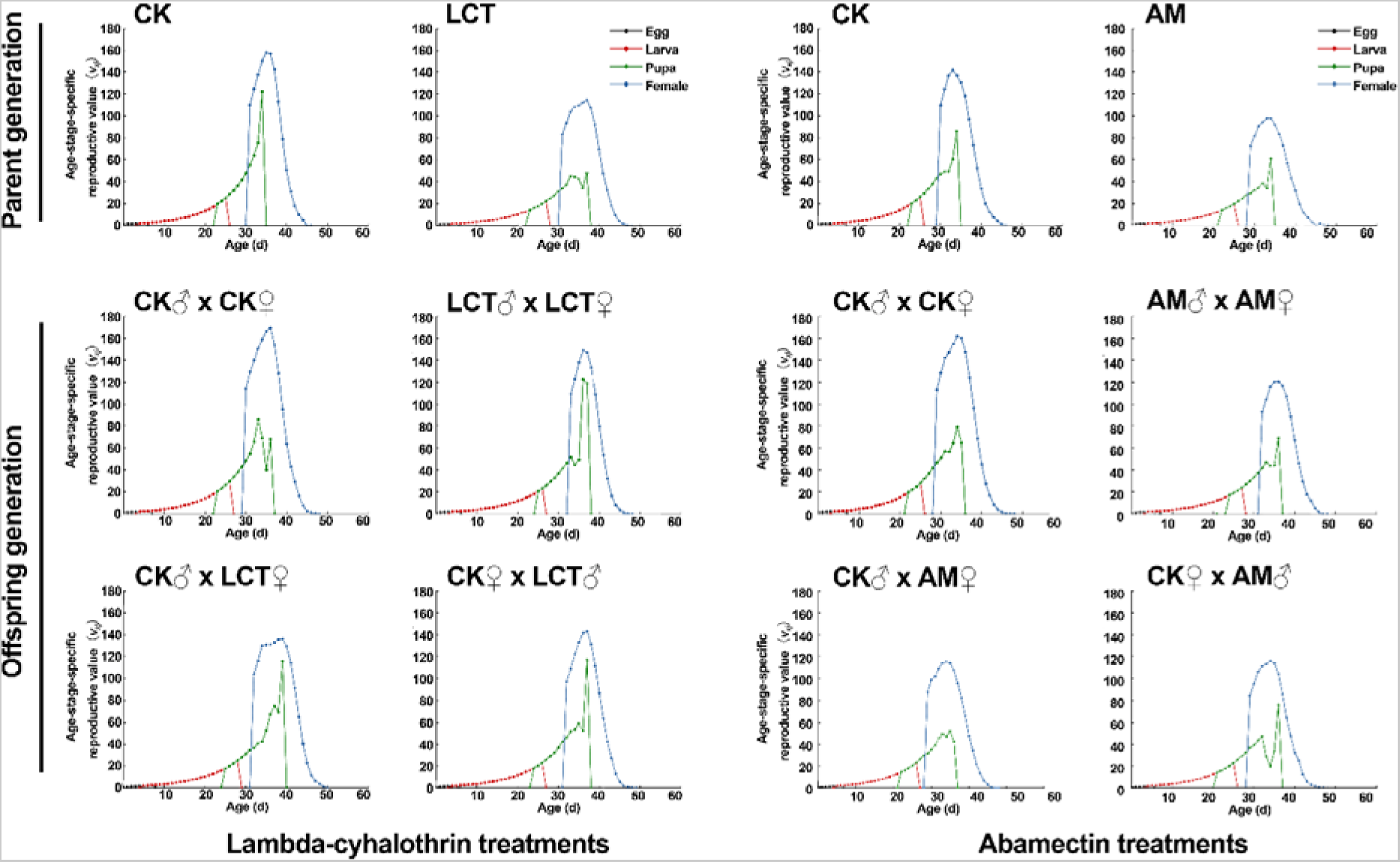
Age-stage-specific reproductive value (*v_xj_*) of *C. pomonella* adult in different treatments. CK, acetone; LCT, lambda-cyhalothrin; AM, abamectin.

### 3.5 Population projection of two successive generations of *C. pomonella*

After beginning with an initial population, the projected population size of *C. pomonella* larvae at 110 days following different treatments is shown in Figure 6 and Figure S4. The total population size of *C. pomonella* larvae after 47 days post exposure (dpe) in the control group was projected to be 3.0-fold larger than the initial population, whereas for LC_30_ of the LCT-treated group, the size was only projected to reach 2.8-fold compared to the initial population. At 85 dpe, the population size of the LCT-treated groups was projected to be 5.0-fold and 4.5-fold in the control and LC_30_, respectively (Figure 6A). In the AM-treated group, the total population sizes at 47 dpe and 85 dpe were projected as 2.8-fold and 4.7-fold, respectively, which were both lower than those in the control group (Figure 6B).

**Figure 6.**
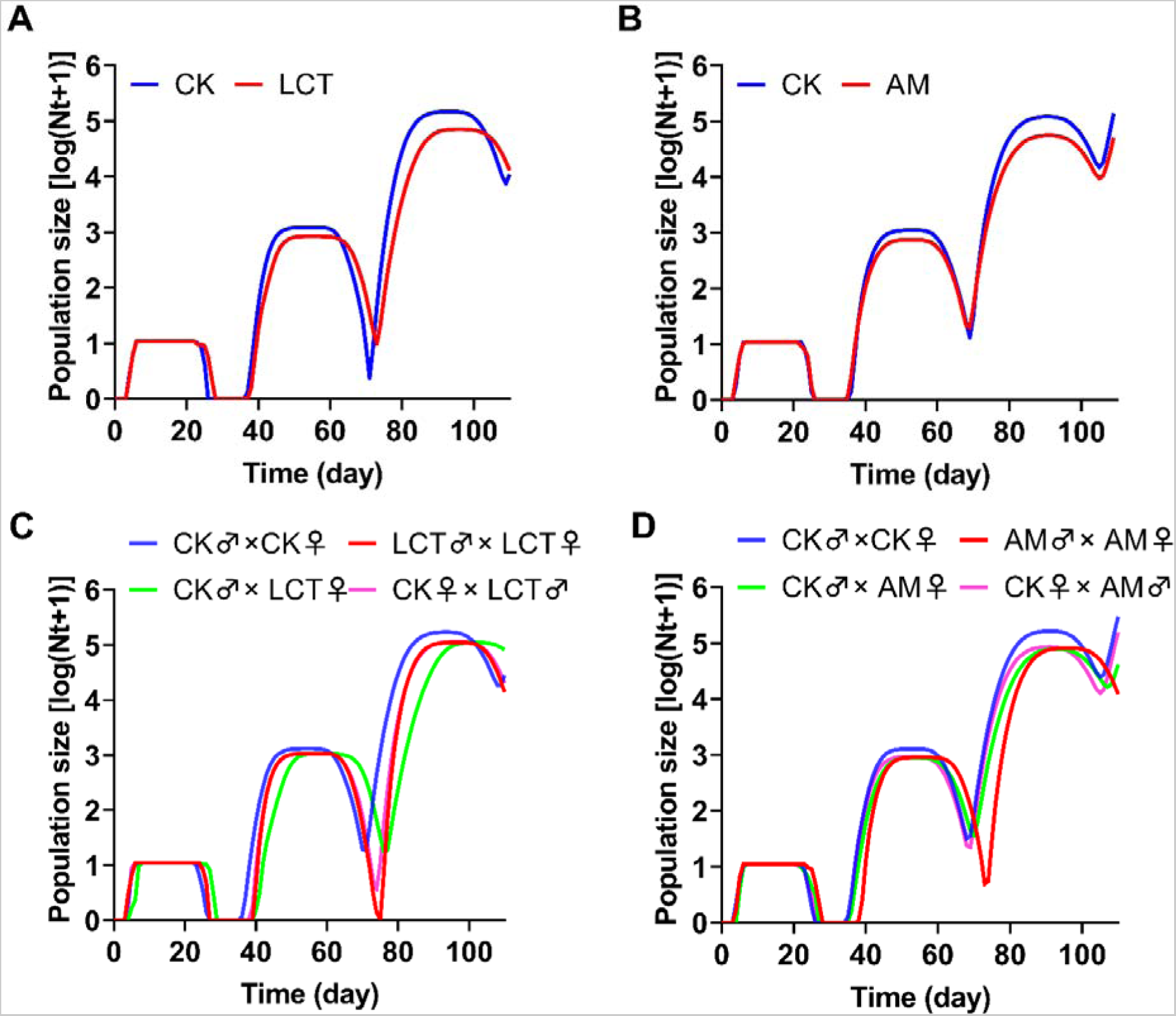
Population projection of *C. pomonella* larvae in different treatments. (A) The estimated population size of *C. pomonella* from an initial population in which neonate larvae were treated with LC_30_ of LCT. (B) The estimated population size of *C. pomonella* from an initial population in which neonate larvae were treated with LC_30_ of AM. (C) The estimated population size of *C. pomonella* from an initial population in which parents (males or females or both) were treated with LC_30_ of LCT. (D) The estimated population size of *C. pomonella* from an initial population in which parents (males or females or both) were treated with LC_30_ of AM. CK, acetone; LCT, lambda-cyhalothrin; AM, abamectin.

Estimated population size was also influenced when beginning with an initial population in which parents (males, females, or both) were treated with LC_30_ of either insecticide. The projected population of all treatments (LCT♂ × LCT♀, CK♂ × LCT♀, CK♀ × LCT♂) was lower when their parents were treated with LCT compared to the control.

Furthermore, in the CK♂ × LCT♀ group, the estimated population size peak was delayed (Figure 6C). The total population sizes at 46 and 84 dpe in the control group were 3.0- and 5.0-fold the initial population, while the value was lower than 3-fold at 46 dpe and 5.0-fold at 86 dpe for all three other treatments (AM♂ × AM♀, CK♂ × AM♀, CK♀ × AM♂) (Figure 6D).

### 3.6 Expression level of *CpVg* gene

The predicted theoretical pI and Mw of the CpVg protein were 8.92 and 18.35 kDa, respectively. Analysis of the protein domain showed that CpVg contained conserved domain Vitellogenin_N (Figure S5). Phylogenetic analysis showed that CpVg was well clustered on branches with vitellogenin proteins among other Lepidoptera insects, suggesting the possibility of close evolutionary relationships between these species (Figure S6).

The mRNA level of *CpVg* was assessed using RT-qPCR (Quantitative reverse transcription PCR) (Figure 7). Results showed that in the parental generation, the *CpVg* transcript was significantly suppressed in female adults emerged from LCT- and AM-treated neonate larvae during the initial and oviposition periods (day 3-5 day after adult emergence), compared with the control. The transcript of *CpVg* in LC_30_ of LCT treated groups were 2.75-, 1.51- and 1.62-fold downregulated at days 3, 4, and 5 after adult emergence, respectively; all significantly lower than those shown in the control (Figure 7A). Similar to LCT treatment, the mRNA level of *CpVg* in AM-treated samples were 5.16-, 4.38-, and 1.95-fold lower than that in the control at day 3, 4, and 5 d after adult emergence, respectively (Figure 7B).

**Figure 7.**
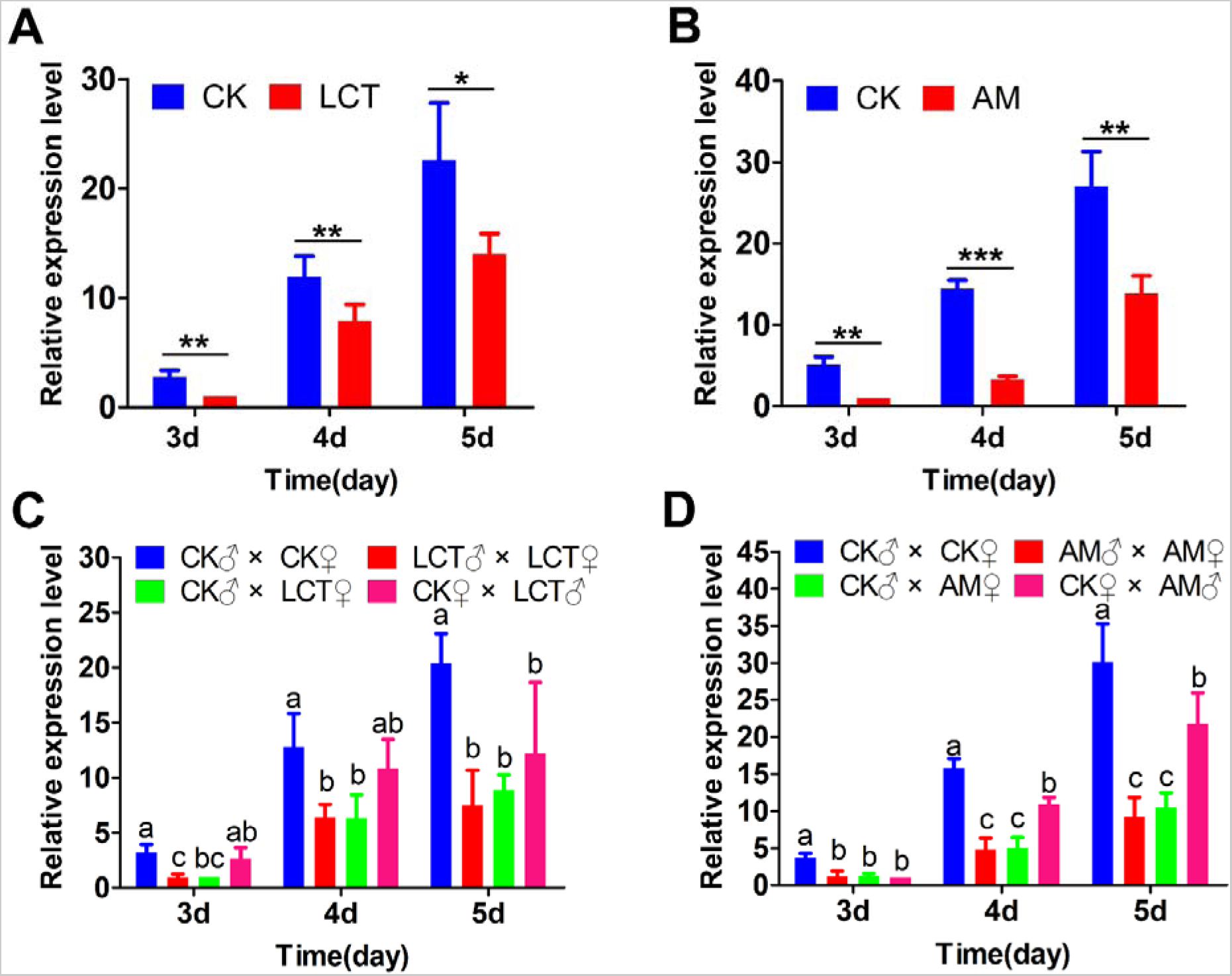
Transgenerational effects of lambda-cyhalothrin (LCT) and abamectin (AM) exposure on the relative expression level of *CpVg* in *C. pomonella* at the 3-5 days after adult emergence. (A) Relative expression of *CpVg* of female moths developed from the larvae treated with LC_30_ of LCT. (B) Relative expression of *CpVg* of female moths developed from the larvae treated with LC_30_ of AM. (C) Relative expression of *CpVg* of offspring whose parents (males or females or both) were treated with LC_30_ of LCT. (D) Relative expression of *CpVg* of offspring whose parents (males or females or both) were treated with LC_30_ of AM. The expression was normalized to transcript levels of reference genes β-*Actin* and *EF-1*α. Three replicates, each replicate with 10 females. Data are presented as mean±SD. Asterisks above represent statistically significant differences by Student’s *t*-test (\**P* ≤0.05; \*\**P* ≤0.01; \*\*\**P* ≤0.001), and letters on the error bars indicate significant differences analyzed by the one-way analysis of variance (ANOVA) with Duncan’s test (*P* < 0.05).

In the offspring generation, the abundance of *CpVg* transcript decreased variably when either or both parents were treated with LCT (Figure 7C) and AM (Figure 7D). Among these treatments, the expression level of *CpVg* in females produced by the mating of insecticide-treated females with non-treated males was not significantly differed from females produced by parents which were both treated with insecticide, and both were significantly lower than the control, and in some cases even lower than shown in the control♀ × insecticide♂ treatment (Figure 7C & 7D).

## 4 Discussion

As a globally important fruit pest, the control of *C. pomonella* primarily depends on chemical insecticides (Reyes et al., 2007; Wang et al., 2019; Hu et al., 2020). Therefore, numerous studies have documented the efficacy and toxicology of insecticides on this species (Yang et al., 2016; Bosch et al., 2018; Wang et al., 2019; Ju et al., 2022). Although neonate larvae are the main targets of insecticides for controlling *C. pomonella* in orchards, the 3rd or 4th instar larvae were mainly assessed, primarily because insecticide application and data analyses were more easily accomplished with larger larvae (Yang et al., 2016; Wang et al., 2019; Ju et al., 2022). However, these results have limited value for guiding the chemical control of *C. pomonella*. Therefore, it is quite meaningful to establish a bioassay method suitable for neonate larvae. In this study, a bioassay method against neonate larvae of a susceptible strain of *C. pomonella* was developed by simulating spraying of insecticides in an orchard. These results showed that the concentration-response lines obtained from this method against neonate larvae showed a meaningful result. Additionally, the variability of mortality in the control was low, therefore the accuracy of the results in this method of insecticide treatment is high. Using this method, the LC_50_ value of LCT obtained in this study (0.22 mg/L) is similar to the result (0.35 mg/L) of Rodríguez et al. (2011) using the insecticide overlaid diet bioassay method against the neonate larvae of a susceptible strain of *C. pomonella*. Furthermore, the LC_50_ value of LCT against neonate larvae at 72 hpe was 0.095 mg/L, which is close to the result (0.02 mg/L) obtained by Mota-Sanchez et al. (2008) at 5 dpe using a susceptible strain. For AM, the LC_90_ value was determined as 0.395 mg/L in the present study and was 0.30 mg/L in previous research (Bosch et al., 2018). These results indicated that the simulated field spray bioassay could be a promising method and might be used effectively in future insecticide resistance monitoring and toxicological research against neonate larvae of *C. pomonella* and other fruit-boring pests.

Pests are inevitably exposed to low and/or sublethal concentrations of insecticides in agro-ecosystems because lethal concentrations of insecticides gradually decrease after initial spraying in fields and orchards (Desneux et al., 2005; 2007). This type of exposure could cause various sublethal effects on pests including negative effects on key life history traits or stimulation of development and/or reproductive capacity (Desneux et al., 2007; Zhen et al., 2018; Li et al. 2018; Liang et al., 2021; Gong et al., 2022; Wu et al., 2022). However, these types of sublethal effects could vary among insect species, insecticide classes and concentrations, and application methods (Müller et al., 2019). Additionally, sensitivity also differs within species depending on developmental stage, sex, and population factors (Gong et al., 2022). Sublethal effects not only occur in exposed individuals but can also occur in offspring (Montaño-Campaz et al., 2022; Gong et al., 2022); however, many studies have focused solely on a single generation (Lai & Su, 2011; Li et al., 2018; Yao et al., 2018). In this study, the transgenerational effects of both maternal and paternal insecticide treatment were taken into consideration. Sublethal concentrations of LCT and AM were found to cause serious negative impacts on the survival rate, body mass, development duration, and reproductive traits of *C. pomonella*. Furthermore, negative consequences in some biological traits might be more obvious in the offspring of insecticide-exposed parents, which were themselves never directly exposed to the insecticides.

Our initial hypothesis was that insecticide treatments impair various performance parameters of the exposed parental generation, and that this can be explained by the toxicity of pyrethroids, even in sublethal concentrations (Desneux et al., 2007; Müller et al., 2017). In this study, the body mass of 3rd instar larvae was found to be significantly reduced, and the survival rate of 4th and 5th instar larvae was seen to be lower than the control. We speculated that this is probably because the larvae underwent the physiological antifeedant effect caused by the insecticides. Although some susceptible individuals survived despite their intestinal absorption system being damaged, insecticide-treated larvae may still have less dietary uptake than the control. Thus, poor nutrition, starvation, or restriction of food during larval stages leads to reduced survival rates of instar larvae (Zhang et al., 2013; Margus, et al., 2019). The weight of female pupae exposed to LCT and male pupae exposed to AM in larval stages was also found to be reduced in comparison to the control. The potential reason is that insecticide-treated larvae must devote more of their energy toward detoxification of insecticides rather than toward growth (Nawaz et al., 2017; Müller et al., 2017; Li et al., 2019). Additional research into the sublethal effects of LCT and AM on the development of *C. pomonella* is essential for elucidating the possible mechanisms.

Sublethal effects of insecticides on the mating behavior and fecundity of adult *C. pomonella* were also observed as lower egg output, reduction of the reproductive period, and decreased production of offspring. Similar results with inhibition of reproduction by LCT or AM were also documented among other insect species including *P. xylostella* (Yin et al., 2008), *Phaedon cochleariae* (Müller et al., 2017) *Scathophaga stercoraria* (van Koppenhagen et al., 2020), and *Phaedon cochleariae* (Wolz et al., 2021a) exposed to pyrethroids or avermectins. Some studies have suggested that decreased fecundity is based on a direct toxic effect of the substances and/or an induced malformation of organs, while others have suggested that exposure to insecticides in larval stages leads to a reduced fat content in females which results in less energy allocation toward egg production (Nawaz et al., 2017; Wolz et al., 2021b). Another suggested possible mechanism is that decreased reproductive output is the result of reduced mating success resulting from exposure to insecticides, and that this may be attributed to insecticide-induced neuron dysfunctions affecting receivers or insecticide effects on chemical cues of the sender (Crawley et al., 2017; Tappert et al., 2017).

It is noteworthy that neither LCT nor AM were observed to significantly affect the population parameters of *C. pomonella*, including the finite rate of increase λ, intrinsic rate of increase *r*, and mean generation time *T*. However, the *R_0_* was shorter in the LCT treated parental generation, and in both generations of AM-treated strains, compared with the control, whereas in comparison to the control, *F* was decreased in both insecticide-treated strains in two successive generations. These results are not in line with results seen in the maize weevil (*S. zeamais*) after being exposed to sublethal doses of pyrethroids insecticide (Guedes et al., 2010), and are different from the results of *S. stercoraria* exposed to sublethal concentrations of ivermectin (van Koppenhagen et al., 2020). Furthermore, Esmaeily et al. (2014) found that *T* was not significantly affected by a low dose of imidacloprid in *Bemisia tabaci* (Esmaeily et al., 2014). These results indicate that the effects of insecticides on insects can vary strongly depending on various factors such as the insecticide chemical family, the insecticide concentration, the endpoints considered, the actual application conditions, the species studied, and the physiological state of the organisms tested (Guo et al., 2013; Chen et al., 2016; Haddi et al., 2016).

Potential transgenerational effects of LC_30_ of LCT and AM on the fitness of *C. pomonella* was also investigated. We found that the larval period was prolonged and the weight of 3rd instar larvae and male pupae was decreased when both larvae were treated with a sublethal concentration of LCT in the parent generation, relative to the control. Meanwhile, the mean generation time (*T*) of offspring was altered when either both parents or the larvae, were exposed to LC_30_ of LCT. These carryover effects may be related to insecticide-induced detoxification mechanisms, leading to trade-offs between developmental and detoxification processes, and resulting in offspring with lower nutrient intake taking a longer time to reach adulthood (Costa et al., 2014; Gibbons et al., 2015; Guedes et al., 2016; Müller et al., 2019; Wolz et al., 2021a). Some previous findings suggested that exposure to sublethal pyrethroids can manifest or induce positive intra- and transgenerational effects, as seen in hormesis effects (Margus et al., 2019; Müller et al., 2019). However, sublethal and transgenerational effects are species-specific and dependent on the life cycle period in which the parents are exposed (Müller et al., 2017; Mauduit et al., 2021).

Generally, avermectin insecticides can have inevitable negative transgenerational effects on offspring biotic performance, which is consistent with previous findings (Galvan et al., 2005; Yin et al., 2008; Costa et al., 2014; Baena-Díaz et al., 2018). In line previous findings that sublethal doses of insecticides in other insects mostly decrease the development rate of immature stages (Gontijo et al., 2014; Nawaz et al., 2017), the present study observed that offspring larval development duration and *T* (mean generation time) tended to be longer in the AM♂ × AM♀ treatment groups than in the control. We also showed that the larvae and pupae of the offspring of the AM-treated parents had lower weights than the offspring of the control parents. This was especially obvious in offspring of exposure treatment mothers. Because mothers can contribute to offspring development through a range of inputs via nutrition, transfer of immune factors, or epigenetic mechanisms, transgenerational effects are most often seen between mother and offspring. This can also be considered as possible causes of the sublethal effects on subsequent generations (Margus et al., 2019; Müller et al., 2019). Additionally, the F (mean fecundity) and R_0_ (net productive rate) of offspring in AM-exposed groups were higher than shown in the control. Some previous studies have indicated that sublethal doses of insecticides might result in the resurgence of target pests (Gong et al., 2022); however, LC_30_ of LCT and AM did not stimulate an increase in the sample population. This result has great significance for IPM program efforts.

Furthermore, *Vg* expression has traditionally been used as an adequate parameter for assessing female fertility (Huang et al., 2016; Yao et al., 2018; Li et al., 2019). In this study, when newly hatched larvae were exposed to sublethal concentrations of LCT or AM, the expression level of *CpVg* was significantly inhibited, and then continuously in 3-5 day post-eclosion female adults. This result was consistent with the observed diminished egg-laying of *C. sinensis* after being exposed to LC_30_ of emamectin benzoate (Yao et al., 2018). Moreover, exposure of 3rd instar larvae of *Chilo suppressalis* to sublethal concentrations of chlorantraniliprole resulted in decreases in both fecundity and Vg expression in female adults (Huang et al., 2016). In contrast, the fecundity and expression level of the *Vg* gene in adult females of *N. lugens* were significantly increased by sublethal concentrations of deltamethrin (Ge et al., 2010). These results indicate that the expression of *Vg* in different insect species varies substantially after sublethal insecticide exposure. Our findings suggest that LCT and AM might influence the reproductive success of *C. pomonella* by modulating the expression of *CpVg*. Moreover, AM decreases the maximal values of *v_xj_* and the total oviposition period and decreases the oviposition peak of offspring. Unsurprisingly, the mRNA expression level of the *CpVg* gene decreased significantly in the offspring generation whose parents were treated with LC_30_ of AM. These results indicated that AM at sublethal concentrations significantly decreased the mRNA expression of the *CpVg* gene in transgenerational adults, along with the observed decrease in fecundity through chronic accumulation. Although the effects of insecticides on transcript levels of *CpVg* may provide an indirect view of the influence of insecticides on the dynamics of ovarian development, the detailed molecular mechanisms of insecticide-induced reproduction, especially hormone related reproduction regulatory mechanisms, require further elucidation.

Our results indicated that sublethal insecticide concentrations may not solely target the exposed generation, but can act transgenerationally and influence unexposed offspring. This is may be explained by a reduced investment in producing offspring after insecticide exposure, insecticide-induced heritable epigenetic changes in genes, or a maternally driven insecticide transfer (Moreau et al., 2012; Müller et al., 2017 & 2019; Wolz et al., 2021a). Importantly, transgenerational effects can also exist if only fathers, and not necessarily mothers, are exposed to insecticides (Baena-Díaz et al., 2018). Transgenerational plasticity, including maternal and paternal effects, is an important tactic used to deal with environmental change, and has been interpreted as cryptic parental care (Baena-Díaz et al., 2018; Mauduit et al., 2021). In the present study, we confirmed the negative consequences of sublethal concentration of two insecticides, LCT and AM, on various parameters over the entire ontogeny of an unexposed offspring generation of *C. pomonella*. Hormesis effects were not observed in *C. pomonella* after insecticide treatment, suggesting that insecticide-induced pest re-emergence will not occur in this species. Additionally, the fecundity and estimated population size was decreased by exposure of neonate larvae to LC_30_ of LCT and AM compared to the control, indicating that the application of these insecticides could provide orchards with longer-term protection from *C. pomonella* damage. However, migration of non-exposure codling moth from untreated apples trees, or other hosts might provide an evolutionary rescue of exposed moths restituting fitness costs in the exposed CM generation, offspring and subsequent generations. Currently, insecticides including LCT have typically been sprayed 8-10 times annually (Wang et al., 2019). Based on these results, we recommend reducing the application frequency of LCT and AM in orchards for managing *C. pomonella*. This may not only reduce the economic costs of pest management, but also promote environmental sustainability and eco-friendliness, including delaying the development of resistance to insecticides in *C. pomonella*.

## 5. Conclusion

In conclusion, this study developed a simulated field spray bioassay method against neonate larvae of *C. pomonella*. Results indicated that LC_30_ of LCT and AM had inevitable intra- and transgenerational sublethal effects on *C. pomonella*. Nevertheless, the situation in orchards may be more complex, including evolutionary rescue by the migration of unexposed individuals from surrounded trees, or other hosts, thus further investigations of the sublethal effects of LCT and AM in the field are recommended. These results may be useful in the development, refining, and implementation of IPM programs against *C. pomonella*.

## Supplementary Information

The online version contains supplementary material available at.

## Acknowledgements

This work was supported by the National Key R&D Program of China (2021YFD1400200).

## Declarations

The authors declare that they have no conflicts of interest.

