## Supplementary Material for "Sublethal and transgenerational effects of lambda-cyhalothrin and abamectin on the development and reproduction of *Cydia pomonella*"

Running title: Sublethal effects of lambda-cyhalothrin and abamectin on *C. pomonella*

[*]Corresponding author: Xue-Qing Yang
College of Plant Protection
Shenyang Agricultural University
Shenyang 110866, Liaoning, China

Orcid ID: 0000-0002-3919-8013

Table S1. Primer sequence used for RT-qPCR.

| primer name | primer sequences (5′→3′) | length (bp) |
| --- | --- | --- |
| *β-Actin*-F | TTGGTATGGGACAGAAGGACTCGTA | 227 |
| *β-Actin*-R | CACGCAGTTCATTGTAGAAGGTGTG | |
| *EF-1α*-F | GGTCCCCTCCAAGCCTCTGT | 157 |
| *EF-1α*-R | CTCGGCAGCTTTGGTGACCT | |
| CPOM07540-F | CCAACATCCAACTATCTTCGTG | 282 |
| CPOM07540-R | TGGCGGTGGGTCCTTTA | |

### Figure legends

**Figure S1.** Vitellogenin mRNA sequence of *C. pomonella*, the specific primer positions were highlighted in green, the PCR product was in red letter.

**Figure S2.** Melt peak (A, A'), amplification plot (B, B'), and melt curve (C, C') of *C. pomonella* vitellogenin gene and reference genes.

**Figure S3.** Sublethal and transgenerational effects of the lambda-cyhalothrin (A) and abamectin (B) on the pupation and emergence and sex ratio of *C. pomonella*. Treatments are given at the bottom, letters on the error bars indicate significant differences analyzed by the one-way analysis of variance (ANOVA) with Duncan's test ($P < 0.05$).

**Figure S4.** Population projection of *C. pomonella* adult in different treatments. (A) Estimated population size of *C. pomonella* from an initial population which neonate larvae treated with $LC_{30}$ of LCT. (B) Estimated population size of *C. pomonella* from an initial population which neonate larvae treated with $LC_{30}$ of AM. (C) Estimated population size of *C. pomonella* from an initial population which parents (males or females or both) were treated with $LC_{30}$ of LCT. (D) Estimated population size of *C. pomonella* from an initial population which parents (males or females or both) were treated with $LC_{30}$ of AM. CK, acetone; LCT, lambda-cyhalothrin; AM, abamectin.

**Figure S5.** Protein domain and secondary structure of *C. pomonella* vitellogenin. (A) The protein of the *C. pomonella* vitellogenin containing conserved domain Vitellogenin_N at interval 45-758 (B) Aignment of the amino acid sequences of *C. pomonella* vitellogenin. The position of α-helices and β-sheets in the amino acid sequence of CpVg were marked as red spiral shape and blue arrow shape, respectively.

**Figure S6.** Phylogenetic analysis of vitellogenin protein from *C. pomonella* (CpVg) and other Lepidoptera insects, a distance neighbor-joining tree was generated using MEGA 7.0. The Vg sequences are obtained from the following GenBank entries: AMD78107.1 for *Chilo suppressalis* (CsVg); QIH04838.1 for *Ostrinia furnacalis* (OfVg); AXY55008.1 for *Maruca vitrata* (MvVg); AEM75020.1 for *Cnaphalocrocis medinalis* (CmVg); XP_048000409.1 for *Leguminivora glycinivorella* (LgVg); XP_014366052.2 for *Papilio machaon* (PmVg); AOH73254.1 for *Spodoptera exigua* (SeVg); ABU68426.1 for *Spodoptera litura* (SlVg); XP_047037412.1 for *Helicoverpa zea* (HzVg); AFV40972.1 for *Helicoverpa armigera* (HaVg); KOB78233.1 for *Operophtera brumata* (ObVg); BAA06397.1 for *Bombyx mori* (BmVg); XP_030021772.1 for *Manduca sexta* (MsVg); BAB32641.1 for *Samia ricini* (SrVg); BAD91195.1 for *Saturnia japonica* (SjVg); ABP63663.1 for *Actias selene* (AsVg).

### Figure S1

```
1     ATGATGTCACTTTGGTATCAGTCTCTCATGCGTCTCGTTCGCGTCGATACATGTACGAACAAATATGAGCGAAATGCACCGGTCGTCGCGTCAGGTCGACATGGCTTAGCCAATCAGAACAACGAAGGCCCCTGGCAGACTGGGAAGCTC
1      M  M  S  L  W  Y  Q  S  L  M  R  L  V  R  V  D  T  C  T  N  K  Y  E  R  N  A  P  V  V  A  S  G  R  H  G  L  A  N  Q  N  N  E  G  P  W  Q  T  G  K  L

151   TACAGCTACGAGGTCCTCTCTAGCACCCTCGCGCGTCTCGAAGAGGGCGCCAGCACCGGCGGCGCCTTCCGGGCCCGCTTCATTGTCCGCGCGCGCGCCCCCGGCCGCCTCCAGGCCAGGCTCGACCAGCCCCAGCACGCCCAGATCCAC
51     Y  S  Y  E  V  L  S  S  T  L  A  R  L  E  E  G  A  S  T  G  G  A  F  R  A  R  F  I  V  R  A  R  A  P  G  R  L  Q  A  R  L  D  Q  P  Q  H  A  Q  I  H

301   CAGGACCTACCTAACACCTGGGCTCTCCCTGTTGACCTCAAATACCAGGACGTTGACAACCTTAACCAAATCTTTGAGATCTATCTCGAGGGTGGCCGTGTTAAGTCACTCAAACTACCAACCTCTGTCTCCCTTCAAAATGAGAATCTG
101    Q  D  L  P  N  T  W  A  L  P  V  D  L  K  Y  Q  D  V  D  N  L  N  Q  I  F  E  I  Y  L  E  G  G  R  V  K  S  L  K  L  P  T  S  V  S  L  Q  N  E  N  L

451   CTCAAAGGCCTCATCAGTGCCCTCCAACTTGACTTATCAGCTCATAGGAACCAGCATTCCGCGCATGATCATCTTGACCAACAAACTCAGCAGGGTCTCTATAGGAAGATGGAGACTGACGTCACTGGTGACTGTGAAACATTGTACGCC
151    L  K  G  L  I  S  A  L  Q  L  D  L  S  A  H  R  N  Q  H  S  A  H  D  H  L  D  Q  Q  T  Q  Q  G  L  Y  R  K  M  E  T  D  V  T  G  D  C  E  T  L  Y  A

601   GTCCTGCCTGTCGCTGCCGAATGGCGCCGAGAACTCCCCAACTTTTCCGATCTCAAGGAAGAGCCTTTCGAAGTCACCAAGAGCAGGGACTACAGACACTGCCACCACAGAGTCGCCTACCACTTTGGAGTGCCCCATGGAGTCCATTGG
201    V  L  P  V  A  A  E  W  R  R  E  L  P  N  F  S  D  L  K  E  E  P  F  E  V  T  K  S  R  D  Y  R  H  C  H  H  R  V  A  Y  H  F  G  V  P  H  G  V  H  W

751   ACCGGACCTCCGGAGAACCCTAAAGACGAGCAGTTCATGAAGCAGAAAATGTACTCTCGCATCCTCGTCGGCAAGCAGGGTCCTATTTACAAGTCTGAGACCACCAGCACAGTTCACGTTAACCCCCATCTTTACGGCAAGCAGAAAGCT
251    T  G  P  P  E  N  P  K  D  E  Q  F  M  K  Q  K  M  Y  S  R  I  L  V  G  K  Q  G  P  I  Y  K  S  E  T  T  S  T  V  H  V  N  P  H  L  Y  G  K  Q  K  A

901   GAGGTACGCAGCAATGTTGTGCTGACTTTAGTCAAGTACGAACAGGACAGTGAGGCCGAATGGCAGGTACCTGAAGGGCGCGTCATCAAGAACCTCCTGTACGCAATGAACTCCAAGCAAATCACTATCCAAGACTCATCGTCTTCCGAA
301    E  V  R  S  N  V  V  L  T  L  V  K  Y  E  Q  D  S  E  A  E  W  Q  V  P  E  G  R  V  I  K  N  L  L  Y  A  M  N  S  K  Q  I  T  I  Q  D  S  S  S  S  E

1051  TCATCGTCATCGGAATCCCACGAGCACCAGCACCATCACTTATACAACCATGAAAGCCAACAGAATGAGGATAACGAGGGCGAAGACAATCAGAACAGAGTGCGCCGATCCTTGAGGTACAAGAAACACGTGACCGTCAACAAGGTGCTC
351    S  S  S  S  E  S  H  E  H  Q  H  H  H  L  Y  N  H  E  S  Q  Q  N  E  D  N  E  G  E  D  N  Q  N  R  V  R  R  S  L  R  Y  K  K  H  V  T  V  N  K  V  L

1201  ATCAAGCGCAGCGATGAAGACAAGTCCTGGGACTCTGACTCGACGTCAGCGTACGTCAACGATGACGTGCCTAACATCAACGAGCCCGCGTACGCGTCCCTATACATGAGCCCCCAGCCCCGCGTGGACAAGAAGCAGAACCCGATGAAC
401    I  K  R  S  D  E  D  K  S  W  D  S  D  S  T  S  A  Y  V  N  D  D  V  P  N  I  N  E  P  A  Y  A  S  L  Y  M  S  P  Q  P  R  V  D  K  K  Q  N  P  M  N

1351  GCCCAGAAACTCGTCCAAGAGATTGCGCAGCTACTTCAGAACCCTAACAACATGCCTAAGAACGACTTCCTCTCCAAGTTTAACATCTTGGTGCGAGTCATTGCCGCTATGAGCTCGGAACAACTCAGCCAGACCAGTCGCAGCATTGAA
451    A  Q  K  L  V  Q  E  I  A  Q  L  L  Q  N  P  N  N  M  P  K  N  D  F  L  S  K  F  N  I  L  V  R  V  I  A  A  M  S  S  E  Q  L  S  Q  T  S  R  S  I  E

1501  GTTGGCAAGTCTTCCAACAACATCCTTAAGTCTGACATGTGGATGGTCCGCGATGCTGTGGCCCAAGCGGGCACCTTACCGGCCTTCCAGCAGATCAAATCTTGGATCGTAAATAAGAAGATCCAAGGAGAGGAGGCCGCTCAGGTCATC
501    V  G  K  S  S  N  N  I  L  K  S  D  M  W  M  V  R  D  A  V  A  Q  A  G  T  L  P  A  F  Q  Q  I  K  S  W  I  V  N  K  K  I  Q  G  E  E  A  A  Q  V  I

1651  GCTTCCCTTCCCGCTACTCTGCGCTACCCAACCAAGGACGTCATGATTCAGTTCTTCAAGCTCGCCATGAGTGATGAGGTTAAGGAGCAGAAGTACTTGAACACCACCACTCTGATTGCCGCCACCAGACTTATCAACCATGCTCAAGTG
551    A  S  L  P  A  T  L  R  Y  P  T  K  D  V  M  I  Q  F  F  K  L  A  M  S  D  E  V  K  E  Q  K  Y  L  N  T  T  T  L  I  A  A  T  R  L  I  N  H  A  Q  V

1801  AACAACGAGACCGCTCACAAATACTATCCCACTCACATGTACGGCCGTCTCGCCAACAAGTACGACAACTTCGTTCTCGAGGAGATCCTGCCGCCCAAAGAGCTGCAGCAAGCGATCACAAACAACGACAACCACAAAGCTCAGGTCTAC
601    N  N  E  T  A  H  K  Y  Y  P  T  H  M  Y  G  R  L  A  N  K  Y  D  N  F  V  L  E  E  I  L  P  P  K  E  L  Q  Q  A  I  T  N  N  D  N  H  K  A  Q  V  Y

1951  ATCAAGGCCATCGGTACTCTCGGCCACCGTAGCATCTTGGATGTCTTTGCTCCGTACTTGGAAGGCAAGGTCCCAACATCCAACTATCTTCGTGCCCAGATGGTGTTGAACCTTGACATGCTAGCTCACGAGAGGGATCACTTCGTTCGT
651    I  K  A  I  G  T  L  G  H  R  S  I  L  D  V  F  A  P  Y  L  E  G  K  V  P  T  S  N  Y  L  R  A  Q  M  V  L  N  L  D  M  L  A  H  E  R  D  H  F  V  R

2101  GCTGCTCTGTACAGTATCCTGAGAAATACGGCTGAGCCCTACGAAGTGCGAGTGTCAGCGATCCACAACATTTTCATGACTCGTCCGACTGTTGCGATGATGCAGGCGATGGCTGAGATGACCCAGAAGGACCCGAGCGTGTACGTCCGC
701    A  A  L  Y  S  I  L  R  N  T  A  E  P  Y  E  V  R  V  S  A  I  H  N  I  F  M  T  R  P  T  V  A  M  M  Q  A  M  A  E  M  T  Q  K  D  P  S  V  Y  V  R

2251  GCTGCGCTTAAGGAAAGCATCCAGTCTGCTGCTGAACTAAAGGACCCACCGCCAGTGCCGCCCAAGCCGCCCAACCTTCCCCTCACCAAGGATGATTTCGGCAGCCTCAACCTAGCCAAGAAATTCGGCGAGTATTTAGGCAGGGATGAA
751    A  A  L  K  E  S  I  Q  S  A  A  E  L  K  D  P  P  P  V  P  P  K  P  P  N  L  P  L  T  K  D  D  F  G  S  L  N  L  A  K  K  F  G  E  Y  L  G  R  D  E

2401  TTTGGGATGATGAACGTCATGTCATACGTCAGCAGCGAGGAGTCCCTGTTCCCGAAGAGCTTGAAGTACTCCTGGAAGAGCAGGTCCTATGGATATCCTAGGATGAACATGATTGCCGCATCGTTCTCAAACTACCGTCAATTTATCGAC
801    F  G  M  M  N  V  M  S  Y  V  S  S  E  E  S  L  F  P  K  S  L  K  Y  S  W  K  S  R  S  Y  G  Y  P  R  M  N  M  I  A  A  S  F  S  N  Y  R  Q  F  I  D

2551  CTGCTGGTGCAAAAGATGACCACGGAGAGACAGACCGCTCCCCAATCGGCCGAGCACAAATTCTCCGCTGAAAAGATCGCTGAAATGCTGAACATCAAGCACGACTCCCAGAAGCCCCTGCTGGCGTCGTTCTTCATCGACTTCATGAAC
851    L  L  V  Q  K  M  T  T  E  R  Q  T  A  P  Q  S  A  E  H  K  F  S  A  E  K  I  A  E  M  L  N  I  K  H  D  S  Q  K  P  L  L  A  S  F  F  I  D  F  M  N

2701  CAACAGAGATATTACAGCCTGACTCAAGAAGATATGGACCGAATTCCAACCCTCTTCGGAGAGTATTTGGCTAATCTGAACAAGGGTTACGACATCCACTACACCAAGGTCTTTAACCAGGCTCAGGTCTCCATCATGTTCCCGCTCGCG
901    Q  Q  R  Y  Y  S  L  T  Q  E  D  M  D  R  I  P  T  L  F  G  E  Y  L  A  N  L  N  K  G  Y  D  I  H  Y  T  K  V  F  N  Q  A  Q  V  S  I  M  F  P  L  A

2851  TCCGGTATGCCTTTCATCTACAAATACAAAGAGCCTACCGTTGTCCACATCCAGGCTAAATCTAAGGGTCAAATCAACATCCCATCTGTCCAGAACCGTGAATATTTCATGAAGCAAGATAATGAGATACAGTTCACATACGCCAGAAAT
951    S  G  M  P  F  I  Y  K  Y  K  E  P  T  V  V  H  I  Q  A  K  S  K  G  Q  I  N  I  P  S  V  Q  N  R  E  Y  F  M  K  Q  D  N  E  I  Q  F  T  Y  A  R  N

3001  ATTGAGGGTAGCGTCGGTTTCTTGAATACGTTGGACAGCCATTTCGTCAGTGCTGGCATTGTCACCAAACACCAGGTTAACCTTGCCTCAAAGGTCCAAATCCAAGCTAAAGCTGGGGAGGTCAAAATCCGTCTTGAACCCCTCCGCCCA
1001   I  E  G  S  V  G  F  L  N  T  L  D  S  H  F  V  S  A  G  I  V  T  K  H  Q  V  N  L  A  S  K  V  Q  I  Q  A  K  A  G  E  V  K  I  R  L  E  P  L  R  P

3151  GAGCAGGAAACCACGATCGCGTACTTTAGCGTCTGGCCTTACACAGCCAATCAGAAGAAGGACTCTCTGACTCCAGTATCCCTCGACCCGACCACGAAAGTCATTACGCGTAGAAACAAAGTGATGTCTGTTGACTACAAGTTCGGACAG
1051   E  Q  E  T  T  I  A  Y  F  S  V  W  P  Y  T  A  N  Q  K  K  D  S  L  T  P  V  S  L  D  P  T  T  K  V  I  T  R  R  N  K  V  M  S  V  D  Y  K  F  G  Q

3301  AGCACTGGCAACCAGTTCCAGCTTCAAGGATATTCGTATTCTAACGATTACAGAAACTACGGAACCATGATGCAGGGCCAGGATCTGCTGTCGAATATCATCTTCGCTCTGAACCAGAGAGACGTTGCCATGACACACTTAAACTTCAGA
1101   S  T  G  N  Q  F  Q  L  Q  G  Y  S  Y  S  N  D  Y  R  N  Y  G  T  M  M  Q  G  Q  D  L  L  S  N  I  I  F  A  L  N  Q  R  D  V  A  M  T  H  L  N  F  R

3451  TACATTGGAAAGCAGTCCCAAAACAAAGCTATTGCTATCACTGTGGCTTATGACACTCTCTACAATCAGAAAGAGTCCGAAGCTCCAGTAGTAGCGTCTGAAATCCAAGACGTTACCCCTATCAGCGCAACCCGCCGTCAAGAGATCGTG
1151   Y  I  G  K  Q  S  Q  N  K  A  I  A  I  T  V  A  Y  D  T  L  Y  N  Q  K  E  S  E  A  P  V  V  A  S  E  I  Q  D  V  T  P  I  S  A  T  R  R  Q  E  I  V

3601  AAACGTGCTGTAGCTGGCATCAAGTACGCCAAAGCTCAGATCATCGATCTGTGCGCCACGTTCGAAGGACCCCAGAAAGCTGAATACATGCTCACCGCCGCCGTCGGCGACAGCAAAGTTGATCCTAAGATCAGATACGCCTTCTTCGCA
1201   K  R  A  V  A  G  I  K  Y  A  K  A  Q  I  I  D  L  C  A  T  F  E  G  P  Q  K  A  E  Y  M  L  T  A  A  V  G  D  S  K  V  D  P  K  I  R  Y  A  F  F  A

3751  GCTAGAAACTCAGCCAAATACGGAACCAACCAGGTCAACTCTGTCGGCAAATTGGACAAGCCTGTCGCATCCGCTCTGAACTTTGTTGAAGCCCTGCAGAAGGAACTGAAGGCCAGCTTCGAAGCGGACTTCAACCTCGGCCAGAAGAAC
1251   A  R  N  S  A  K  Y  G  T  N  Q  V  N  S  V  G  K  L  D  K  P  V  A  S  A  L  N  F  V  E  A  L  Q  K  E  L  K  A  S  F  E  A  D  F  N  L  G  Q  K  N

3901  GGAAATATTCAGATCAAGGGATCAGCTGAGCGCACCAAAAAAGCCACTGAAGAGATCCAACAGAGCCCCTGGGCCAAGCAGAGCGTCCAAGAAATCGCCCGCAACAACTACTACCAGGAGGCCAGCAGAATGATGATCGAAATGGCGTAC
1301   G  N  I  Q  I  K  G  S  A  E  R  T  K  K  A  T  E  E  I  Q  Q  S  P  W  A  K  Q  S  V  Q  E  I  A  R  N  N  Y  Y  Q  E  A  S  R  M  M  I  E  M  A  Y

4051  GCTCCCACTCACGTTCTGGCTTCGGTATCCTACAAAGATTTAAGCCCTTCCGTCATCAACATGACCTACCACGCTTACAAGGCACTGACCACCTACGGTTACTGGTATTCTGATATCAACGTAATGAAAAAATCTGCTGAGGGTAAACTC
1351   A  P  T  H  V  L  A  S  V  S  Y  K  D  L  S  P  S  V  I  N  M  T  Y  H  A  Y  K  A  L  T  T  Y  G  Y  W  Y  S  D  I  N  V  M  K  K  S  A  E  G  K  L

4201  GATTTCGATGCGGAGGCGTCCTACGTGGATAACTACCTGAACTTGAACATGATCTCACGATATGGCCATCTTAAGATGAACAACGTACCAATCCCCAAGACCAGCGCTGCTGCTATGACGTTCTACGCGCCGCTGAATGCTTATGAGTGC
1401   D  F  D  A  E  A  S  Y  V  D  N  Y  L  N  L  N  M  I  S  R  Y  G  H  L  K  M  N  N  V  P  I  P  K  T  S  A  A  A  M  T  F  Y  A  P  L  N  A  Y  E  C

4351  GTCCTAAATCGATTCACCAGGCAACAATACCTGCCCTACTGCAGCGTTGACGGTTCAAAGATCAGGACGTTCAGCAACCGAAGCTACGACTACACCCTAACCAGTTCCTGGCACGTGGTGGTCCAGGACAACGCCAAGTACGCTGGACAG
1451   V  L  N  R  F  T  R  Q  Q  Y  L  P  Y  C  S  V  D  G  S  K  I  R  T  F  S  N  R  S  Y  D  Y  T  L  T  S  S  W  H  V  V  V  Q  D  N  A  K  Y  A  G  Q

4501  CAAGGAGTGAGCAACGAGCAGCTCGTCATCCTGGCTAGGAGGCCTGAGGAGAAGAAGCAGGAGATCTATGTTTCGTTCAAATCTGACACCGGCAATCACTTGGAAATGGAAGTCCAGCCTGCAGCGAGTCTCAACTTAAAGGAAGGATTG
1501   Q  G  V  S  N  E  Q  L  V  I  L  A  R  R  P  E  E  K  K  Q  E  I  Y  V  S  F  K  S  D  T  G  N  H  L  E  M  E  V  Q  P  A  A  S  L  N  L  K  E  G  L

4651  AAAGTAACCACCAATGCCAAGAAAGTCTCCGACGGTGACTTCACCATGTATTGGGATGAAGCCAAGGACGCCCCCTCTTCTGTATTACTACAATACCGATGGTGCGTGGATGTTCGTGATTGGAGACGATCGTCTCCGCGGCATTTACGA
1551   K  V  T  T  N  A  K  K  V  S  D  G  D  F  T  M  Y  W  D  E  A  K  D  A  P  S  S  V  L  L  Q  Y  R  W  C  V  D  V  R  D  W  R  R  S  S  P  R  H  L  R

4801  TGGCCAGAGACTCGTGATCTGGGCGCGCAATAA
1601   W  P  E  T  R  D  L  G  A  Q  *
```

**Figure S2**

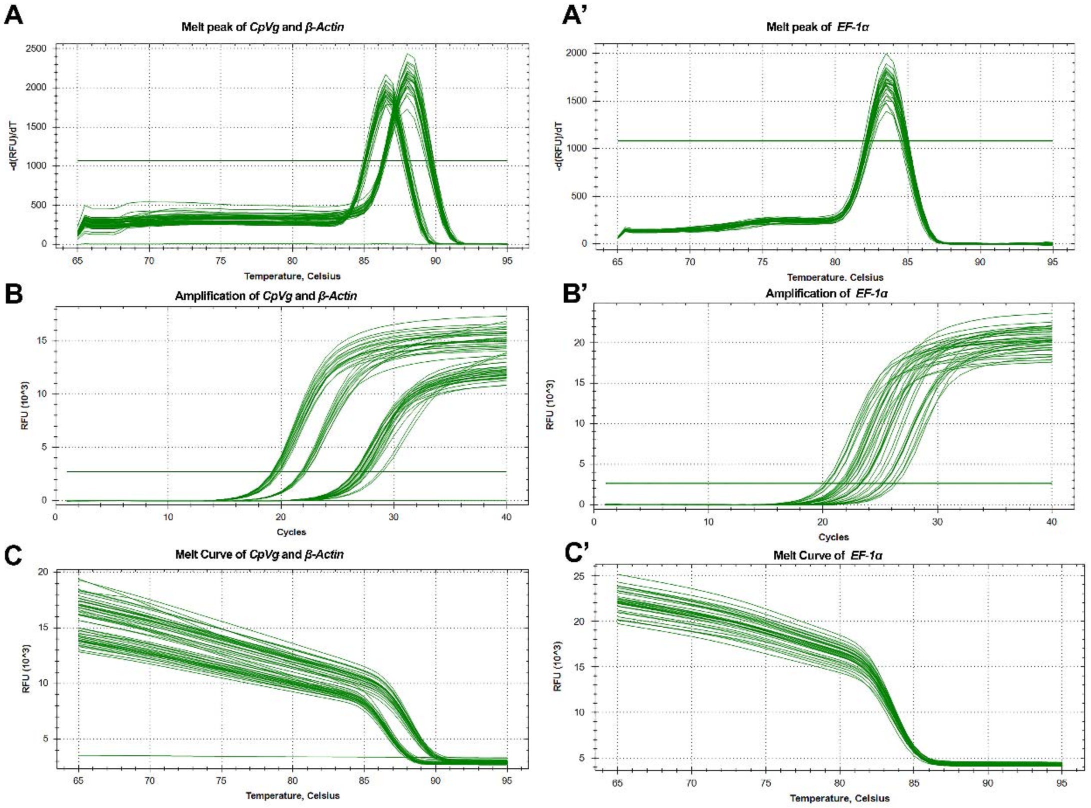

A
Melt peak of CpVg and β-Actin
-d(RFU)/dT
Temperature, Celsius
A’
Melt peak of EF-1α
-d(RFU)/dT
Temperature, Celsius
B
Amplification of CpVg and β-Actin
RFU (10^3)
Cycles
B’
Amplification of EF-1α
RFU (10^3)
Cycles
C
Melt Curve of CpVg and β-Actin
RFU (10^3)
Temperature, Celsius
C’
Melt Curve of EF-1α
RFU (10^3)
Temperature, Celsius

**Figure S3**

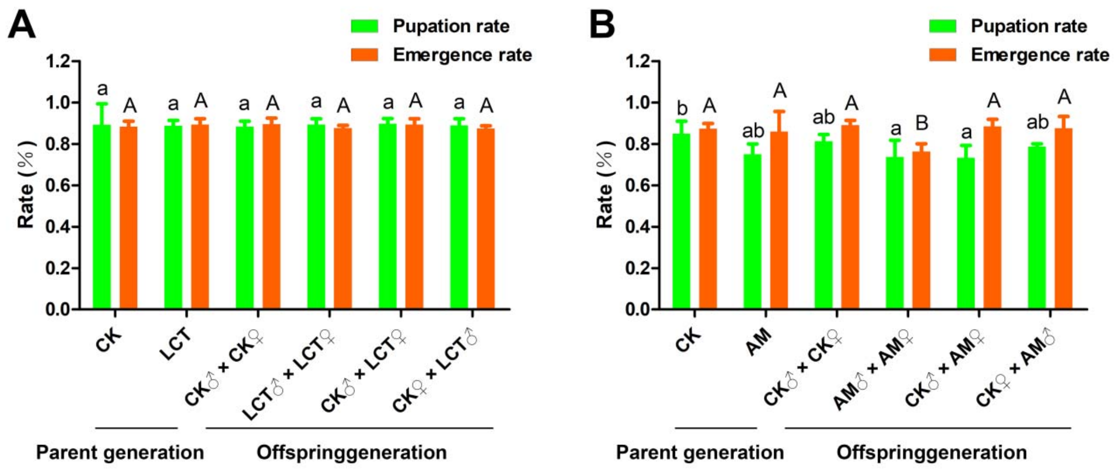

A
Pupation rate
Emergence rate
Rate (%)
1.2
1.0
0.8
0.6
0.4
0.2
0.0
a A a A a A a A a A a A
CK
LCT
CK♂ × CK♀
LCT♂ × LCT♀
CK♂ × LCT♀
CK♀ × LCT♂
Parent generation
Offspringgeneration
B
Pupation rate
Emergence rate
Rate (%)
1.2
1.0
0.8
0.6
0.4
0.2
0.0
b A ab A ab A a B a A ab A
CK
AM
CK♂ × CK♀
AM♂ × AM♀
CK♂ × AM♀
CK♀ × AM♂
Parent generation
Offspringgeneration

Figure S4

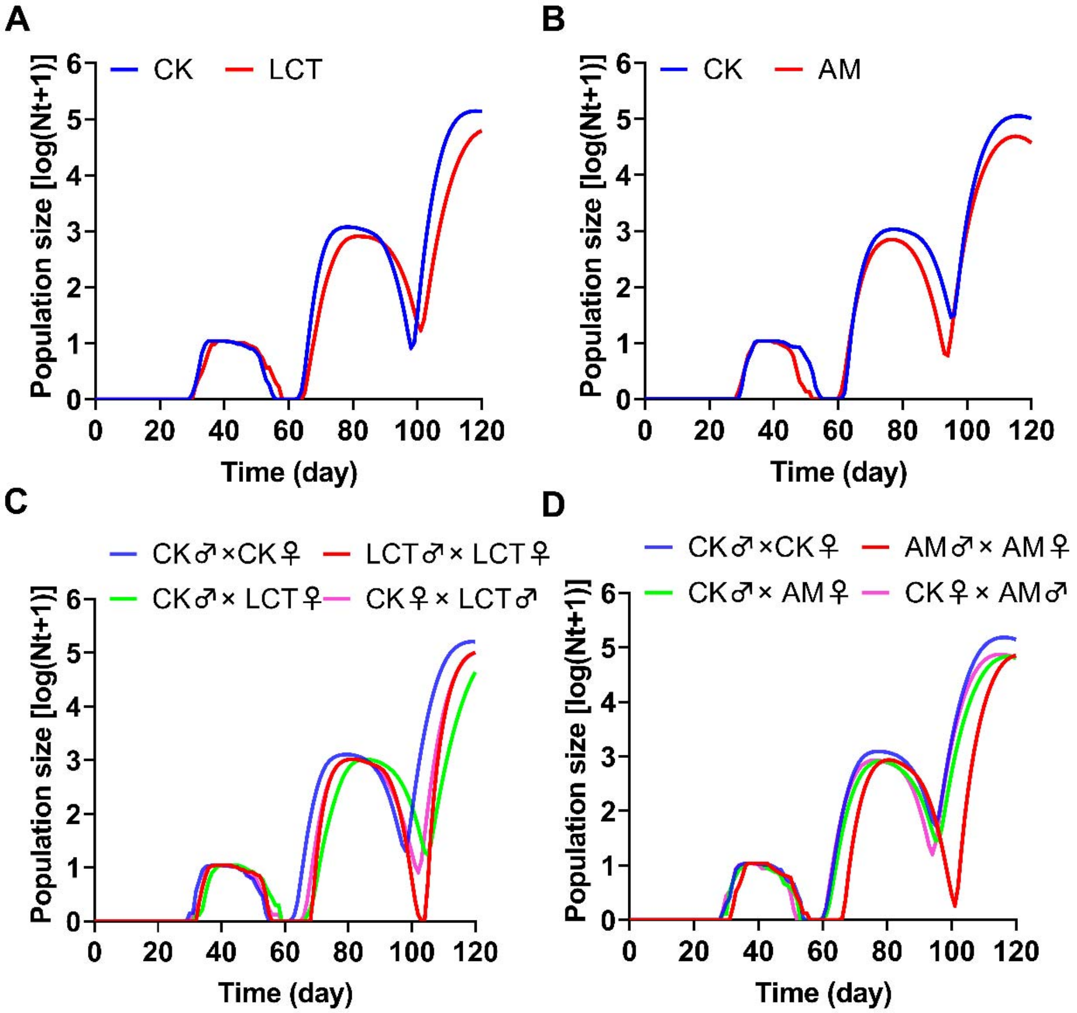

A
CK
LCT
Population size [log(Nt+1)]
Time (day)
B
CK
AM
Population size [log(Nt+1)]
Time (day)
C
CK♂×CK♀
LCT♂× LCT♀
CK♂× LCT♀
CK♀× LCT♂
Population size [log(Nt+1)]
Time (day)
D
CK♂×CK♀
AM♂× AM♀
CK♂× AM♀
CK♀× AM♂
Population size [log(Nt+1)]
Time (day)

### Figure S5

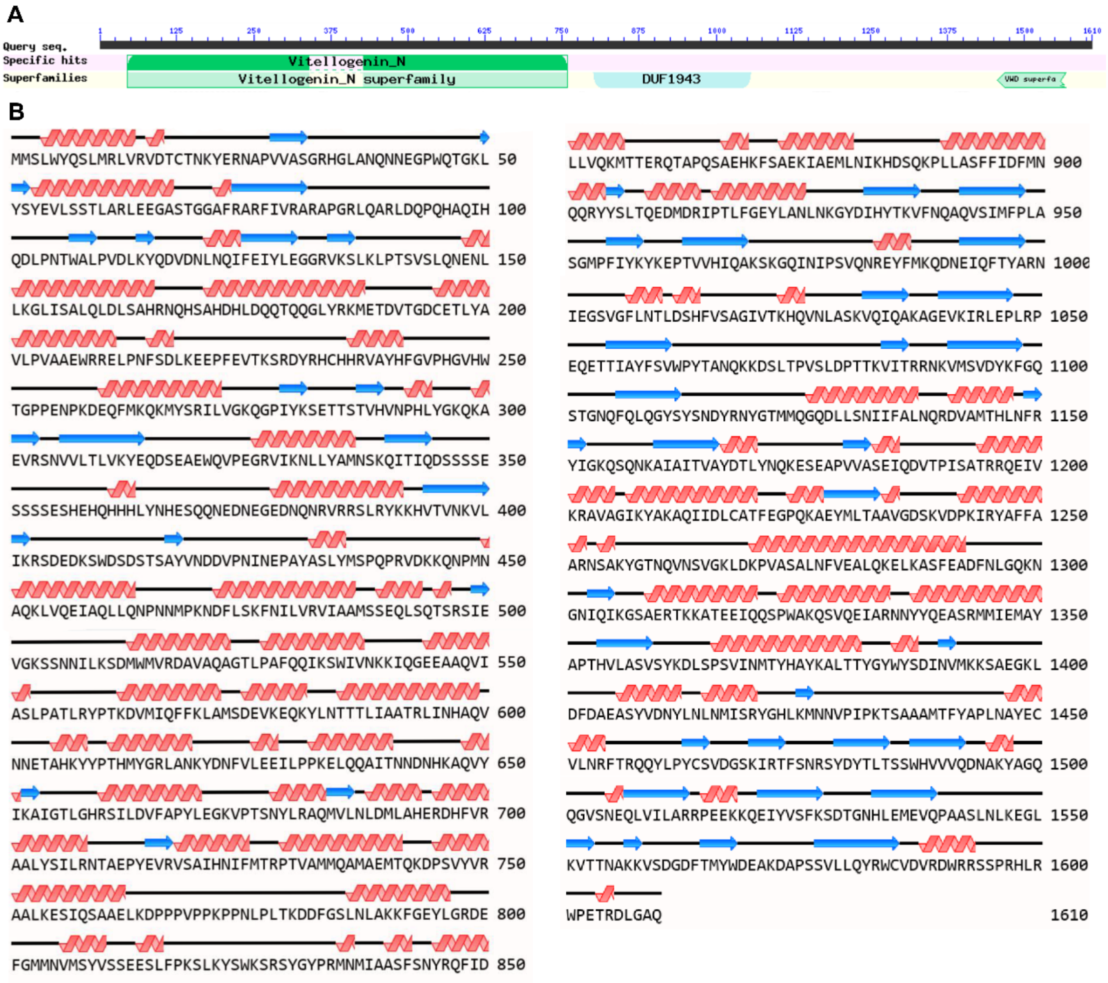
A
1 125 250 375 500 625 750 875 1000 1125 1250 1375 1500 1610
Query seq.
Specific hits
Superfamilies
Vitellogenin_N
Vitellogenin_N superfamily
DUF1943
VWD superfa
B
MMSLWYQSLMRLVRVDTCTNKYERNAPVVASGRHGLANQNNEGPWQTGKL 50
YSYEVLSSTLARLEEGASTGGAFRARFIVRARAPGRLQARLDQPQHAQIH 100
QDLPNTWALPVDLKYQDVDNLNQIFEIYLEGGRVKSLKLPTSVSLQNENL 150
LKGLISALQLDLSAHRNQHSAHDHLDQQTQQGLYRKMETDVTGDCETLYA 200
VLPVAAEWRRELPNFSDLKEEPFEVTKSRDYRHCHHRVAYHFGVPHGVHW 250
TGPPENPKDEQFMKQKMYSRILVGKQGPIYKSETTSTVHVNPHLYGKQKA 300
EVRSNVVLTLVKYEQDSEAEWQVPEGRVIKNLLYAMNSKQITIQDSSSSE 350
SSSSESHEHQHHHLYNHESQQNEDNEGEDNQNRVRRSLRYKKHVTVNKVL 400
IKRSDEDKSWDSDSTSAYVNDDVPNINEPAYASLYMSPQPRVDKKQNPMN 450
AQKLVQEIAQLLQNPNNMPKNDFLSKFNILVRVIAAMSSEQLSQTSRSIE 500
VGKSSNNILKSDMWMVRDAVAQAGTLPAFQQIKSWIVNKKIQGEEAAQVI 550
ASLPATLRYPTKDVMIQFFKLAMSDEVKEQKYLNTTTLIAATRLINHAQV 600
NNETAHKYYPTHMYGRLANKYDNFVLEEILPPKELQQAITNNDNHKAQVY 650
IKAIGTLGHRSILDVFAPYLEGKVPTSNYLRAQMVLNLDMLAHERDHFVR 700
AALYSILRNTAEPYEVRVSAIHNIFMTRPTVAMMQAMAEMTQKDPSVYVR 750
AALKESIQSAAELKDPPPVPPKPPNLPLTKDDFGSLNLAKKFGEYLGRDE 800
FGMMNVMSYVSSEESLFPKSLKYSWKSRSYGYPRMNMIAASFSNYRQFID 850
LLVQKMTTERQTAPQSAEHKFSAEKIAEMLNIKHDSQKPLLASFFIDFMN 900
QQRYYSLTQEDMDRIPTLFGEYLANLNKGYDIHYTKVFNQAQVSIMFPLA 950
SGMPFIYKYKEPTVVHIQAKSKGQINIPSVQNREYFMKQDNEIQFTYARN 1000
IEGSVGFLNTLDSHFVSAGIVTKHQVNLASKVQIQAKAGEVKIRLEPLRP 1050
EQETTIAYFSVWPYTANQKKDSLTPVSLDPTTKVITRRNKVMSVDYKFGQ 1100
STGNQFQLQGYSYSNDYRNYGTMMQGQDLLSNIIFALNQRDVAMTHLNFR 1150
YIGKQSQNKAIAITVAYDTLYNQKESEAPVVASEIQDVTPISATRRQEIV 1200
KRAVAGIKYAKAQIIDLCATFEGPQKAEYMLTAAVGDSKVDPKIRYAFFA 1250
ARNSAKYGTNQVNSVGKLDKPVASALNFVEALQKELKASFEADFNLGQKN 1300
GNIQIKGSAERTKKATEEIQQSPWAKQSVQEIARNNYYQEASRMMIEMAY 1350
APTHVLASVSYKDLSPSVINMTYHAYKALTTYGYWYSDINVMKKSAEGKL 1400
DFDAEASYVDNYLNLNMISRYGHLKMNNVPIPKTSAAAMTFYAPLNAYEC 1450
VLNRFTRQQYLPYCSVDGSKIRTFSNRSYDYTLTSSWHVVVQDNAKYAGQ 1500
QGVSNEQLVILARRPEEKKQEIYVSFKSDTGNHLEMEVQPAASLNLKEGL 1550
KVTTNAKKVSDGDFTMYWDEAKDAPSSVLLQYRWCVDVRDWRRSSPRHLR 1600
WPETRDLGAQ 1610

**Figure S6**

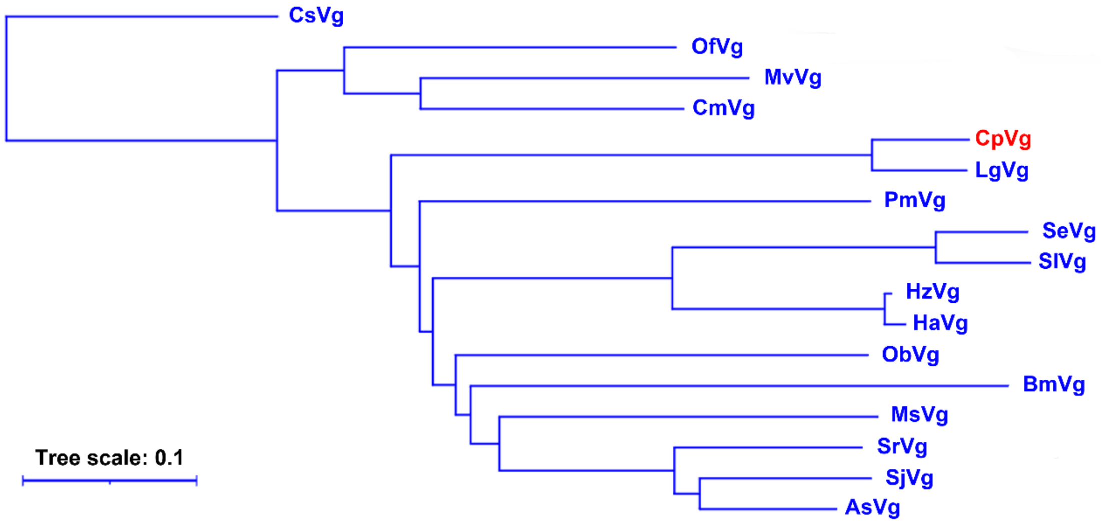
CsVg
OfVg
MvVg
CmVg
CpVg
LgVg
PmVg
SeVg
SlVg
HzVg
HaVg
ObVg
BmVg
MsVg
SrVg
SjVg
AsVg
Tree scale: 0.1
